# Inhibition of cortico-amygdala projections underlies affective bias modification by psilocybin

**DOI:** 10.64898/2026.03.02.709133

**Authors:** Matthew D. B. Claydon, Justyna K. Hinchcliffe, Julia Bartlett, Caroline T. Golden, Christopher W. Thomas, Gary Gilmour, Jack R. Mellor, Zuner A. Bortolotto, Emma S. J. Robinson

## Abstract

Psilocybin, a serotonergic psychedelic, can produce rapid and enduring antidepressant effects in patients with major depressive disorder (MDD)[1, 2], yet the neural mechanisms underlying these effects remain unclear. Negative affective biases are an important neuropsychological mechanism central to the development and perpetuation of MDD[3]. Using a translational rodent model, we previously demonstrated that psilocybin modulates negative affective biases which, we hypothesize, contribute to its antidepressant effects[4].

Here, we identify the prelimbic subregion (PrL) of the rat medial prefrontal cortex (mPFC) as a key locus for the modulation of affective biases by psilocin, the active metabolite of psilocybin, and reveal a cell-type-specific bidirectional regulation of synaptic transmission. Psilocin selectively suppressed excitatory synaptic input to cortico-amygdala (CA) projection neurons, but enhanced excitatory transmission to other, putatively cortico-cortical, targets. Interestingly, suppression of the excitatory input to CA cells by psilocin, and modulation of affective biases by psilocybin, were both dependent on 5HT_1A_ and 5HT_2A_ receptor signaling.

Consistent with the long-term therapeutic effects of rapidly acting antidepressants[1, 2, 4, 5], psilocin produced sustained changes to affective biases evident 24 hours after PrL infusion. In parallel, the suppressed excitatory transmission shifted to enhanced inhibitory synaptic input selectively in CA cells. Finally, chemogenetic inhibition of CA neurons in PrL recapitulated both the acute and sustained modulation of negative affective biases by psilocybin, as well as positively biasing new reward memories.

Together, these findings identify modulation of the PrL cortico-amygdala circuit as a key substrate for affective bias modification by psilocybin, an effect which could explain its rapid and sustained antidepressant actions.

## Introduction

Major depressive disorder (MDD) is a leading cause of disability worldwide, yet treatment outcomes remain limited[6, 7]. This is, in part, due to the complexity and heterogeneity of symptoms, as well as a lack of understanding of the underlying neurobiological mechanisms[8–10]. Conventional treatment options like selective serotonin reuptake inhibitors (SSRIs) have limited efficacy and a delayed onset of action. Indeed, an estimated 30-40% of patients are failed by current treatments, and side-effects associated with these long-standing treatment options can be significant[11, 12]. Recently, rapid acting antidepressants (RAADs), like the NMDA receptor antagonist, ketamine, and the serotonergic psychedelic, psilocybin, have emerged as potentially efficacious options in treatment-resistant patients[1, 2, 5, 13]. In some clinical trials, psilocybin promotes rapid and sustained improvements in a proportion of the participants’ mood following a single dose[1], and neuroplastic mechanisms have been postulated to be an important driver of the durable effects. However, an understanding of the precise nature of these mechanisms remains limited.

Mechanistic studies of psychiatric disorders are inherently challenging, and while limited, animal models remain essential for bridging neurobiology and behavior. Traditional rodent assays of antidepressant efficacy rely on behavioral readouts that have limited translational validity, particularly when investigating underlying mechanisms[14, 15]. In contrast, affective bias paradigms, tasks that quantify how emotional states shape cognition, may provide a tractable framework linking rodent and human behavior[16, 17]. Negative affective biases are a core neuropsychological mechanism central to the low mood and anhedonia that are characteristic of MDD, and their alleviation may represent a key substrate of antidepressant response[3]. Indeed, a single low dose (0.3 mg/kg) of psilocybin attenuates a memory specific, negative affective bias and facilitates re-learning with a more positive affective valence 24hrs post-treatment[4]. Acute psilocybin also generates positive affective biases associated with new experiences, similar to the effects seen with conventional antidepressants [4, 16, 18]. Yet, the neurobiological mechanisms through which psilocybin produces these lasting neuropsychological effects remain unknown.

At the molecular level, psilocybin acts via its active metabolite, psilocin, which agonizes various serotonin receptors[14]. Psychedelic, plasticity-promoting, and therapeutic effects are thought to be mediated primarily via the Gq coupled 5HT_2A_ receptor[19–21], although the Gi coupled 5HT_1A_ receptor has also been linked to antidepressant mechanisms[22], and the unique poly-pharmacology of psilocybin may be crucial to its actions[23–25]. 5HT receptor subtypes are differentially distributed across the central nervous system[26]; however, converging evidence suggests that the medial prefrontal cortex (mPFC) is a key site of modulation by psilocybin and other psychedelic drugs[19, 20, 27]. Indeed, this region plays an important role in regulating emotional processing and cognitive flexibility[28]. Human neuroimaging studies reveal that psilocybin alters prefrontal activity and functional connectivity with limbic structures such as the amygdala, correlating with therapeutic outcomes[29]. Preclinical studies indicate that psilocybin induces structural plasticity in cortical pyramidal neurons[19, 20], but the specific cell types and circuits altered in the modulation of affective processing remain unclear.

Here, we combine behavioral, electrophysiological, and chemogenetic approaches to identify the neural substrates of psilocybin-mediated affective bias modification. We identify prelimbic mPFC (PrL) as a key locus for these actions. Psilocin bidirectionally regulates excitatory transmission within this region, selectively suppressing transmission onto cortico-amygdala (CA) projection neurons through coordinated actions at 5-HT_2A_ and 5-HT_1A_ receptors. Sustained modulation of affective biases was accompanied by enhanced inhibitory input onto CA neurons 24 hours after drug exposure. Finally, chemogenetic inhibition of CA neurons reproduced both the acute and 24h post-treatment behavioral effects of psilocybin. Together, these findings identify the CA circuit as a key substrate for psilocybin’s modulation of affective bias, providing mechanistic insight into how this compound produces rapid and enduring antidepressant effects.

## Results

### Psilocybin modulates negative affective biases by actions in prelimbic medial prefrontal cortex

In the rodent affective bias task (ABT), digging substrates provide cues which are predictive of a food reward. To induce an affective bias, each rat learns two independent substrate-reward associations over four pairing sessions with treatment/control, substrate and order of presentation fully counter-balanced (Table S2). The value of the food reward is kept constant throughout and the affective bias generated quantified using a choice test where the animals are presented with the two previous reward-associated digging substrates at the same time and their choices over 30 randomly reinforced trials recorded. We have previously shown that the negatively biased memory generated by the benzodiazepine inverse agonist, FG-7142, can be acutely attenuated by low systemic doses of psilocybin (0.3 mg/kg (intraperitoneal (IP))[4].

To investigate modulation of a negatively biased memory by psilocybin’s actions in the PrL specifically, we first generated a negative affective bias using FG-7142[16] (3mg/kg sub-cutaneous (SC)) and then infused psilocin (psilocybin’s active metabolite) or vehicle via an intra-PrL cannula 5 minutes before the choice test (Figure 1A).

**Figure 1.**
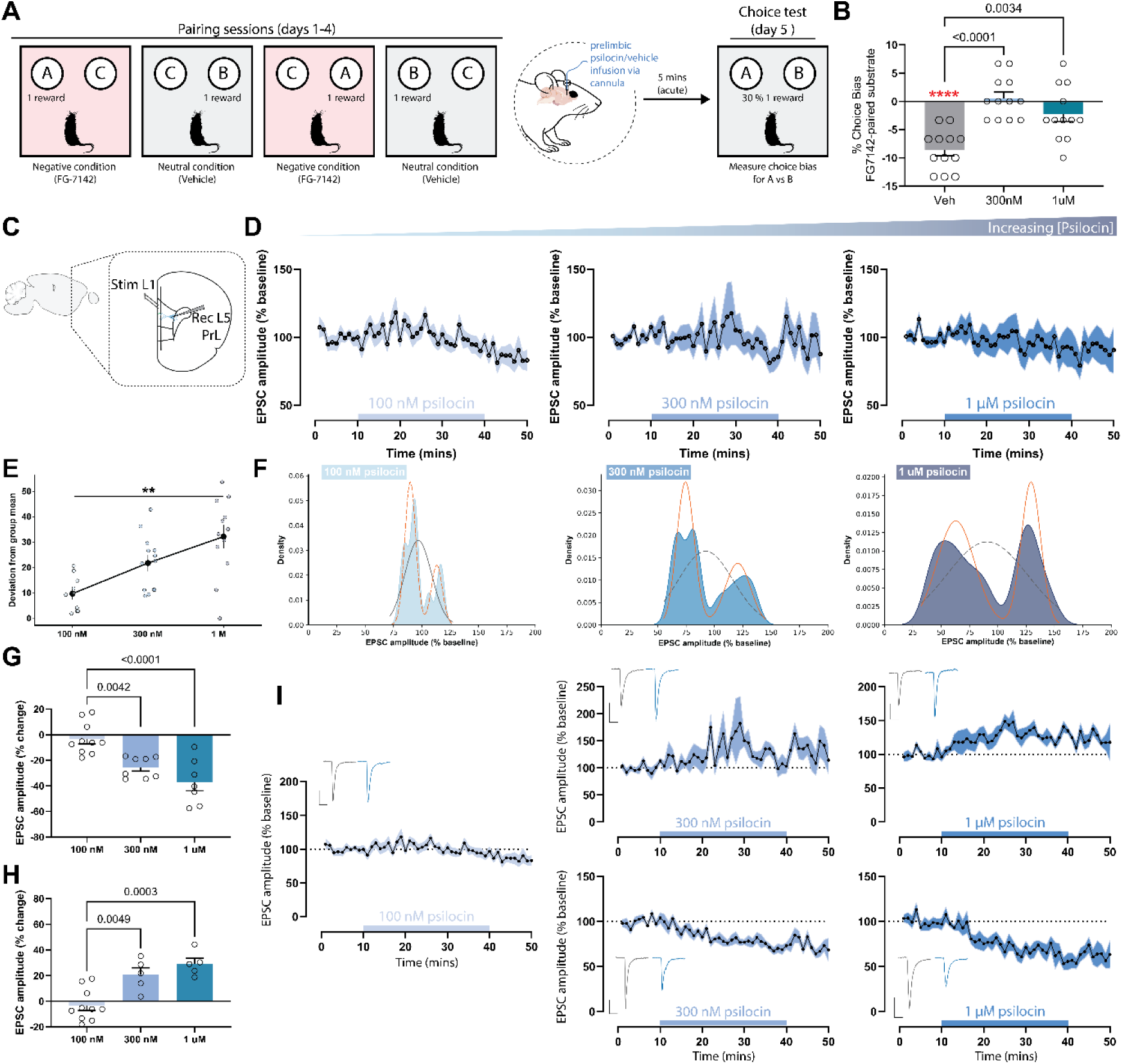
Psilocybin modulates negative affective biases by acting on prelimbic mPFC with evidence of bimodal effects involving subpopulations of layer 5 pyramidal neurons. **A)** Overview of the affective bias test (ABT) protocol used to generate a negative affective bias and investigate the acute effects of psilocin infusion. Animals were treated with FG-7142 (3mg/kg SC) to induce a negatively biased memory. Affective biases were quantified using a choice test with psilocin or vehicle administered by infusion 5 minutes before testing to investigate the acute effects on retrieval. **B)** Choice biases in male rats (n = 12 per group) following prelimbic infusion of vehicle, 300 nM psilocin, or 1 µM psilocin. Data were analyzed using one-sample t-tests against a null hypothesis mean of 0% choice bias (Veh: t₁₁ = 8.258, ****P < 0.0001) and a repeated-measures ANOVA with Dunnett’s post hoc comparisons (main effect of treatment: F_2, 22_ = 13.73, P = 0.0001). **C)** Schematic of whole-cell recordings from layer 5 prelimbic pyramidal neurons held at –70 mV while EPSCs were evoked by layer 1 stimulation. **D)** Time series data showing EPSC amplitudes recorded before, during, and after varying concentrations of psilocin wash-in: 100 nM (n = 10), 300 nM (n = 12), or 1 µM (n = 12). **E)** Variance in EPSC amplitude during the final 10 minutes of wash-in was related to psilocin concentration as determined by ordinary least squares regression (β = 21.7 ± 5.4 (SE), t₃₃ = 4.1, P = 0.0003) and Levene’s test (F = 6.49, **P = 0.004). **F)** Gaussian mixture modelling of EPSC amplitudes during the final 10 minutes of wash-in showing model fits for single- and two-component Gaussian distributions. Model comparison using BIC indicated a better fit for a two-component model at 300 nM (BIC₂ = 123.62; BIC₁ = 124.93) and 1 µM (BIC₂ = 123.49; BIC₁ = 124.70) psilocin, and a better fit for a single Gaussian at 100 nM (BIC₁ = 82.25; BIC₂ = 83.61). **G-H)** EPSC amplitudes binned according to mixture-model components, showing groups of neurons exhibiting potentiated or depressed responses across psilocin concentrations. Potentiated cells were analyzed using a one-way ANOVA (main effect of drug: F_2, 17_ = 14.64, P = 0.0002), and depressed cells using a one-way ANOVA (main effect of drug: F_2, 22_ = 13.39, P = 0.0002), followed by Tukey’s post hoc comparisons. **I)** Binned time-series data corresponding to the potentiated and depressed neuronal subpopulations. Representative traces averaged across 10 minute time bins are baseline (grey) and final 10 minutes of psilocin (blue), scale bars are x = 50 ms, y = 50 pA. Data are mean ± SEM.

During the choice test, rats receiving vehicle infusions made fewer choices for the FG-paired substrate, consistent with a negatively biased memory (one-sample t-test, ****P < 0.0001, Figure 1B). Infusion of both 300 nM and 1 μM of psilocin into PrL attenuated the negative affective bias (main effect of treatment: F_2, 22_ = 13.73, P = 0.0001, RM-ANOVA with Dunnett’s; Figure 1B), replicating the effects of systemic low-dose psilocybin, and identifying PrL as a key locus for the affective bias modification induced by psilocybin. To gain mechanistic insight, we next sought to investigate the effects of psilocin on layer V pyramidal output neurons of the PrL.

Excitatory post-synaptic currents (EPSCs) in L5 pyramidal cells were elicited in ex-vivo slice preparations by stimulation of the layer I fiber tracts (Figure 1C). Stable EPSC baselines were recorded and psilocin (100 nM, 300 nM, and 1 μM) was bath applied to the slices. On average there was no change to EPSC amplitude (Figure 1D), however, we observed a concentration-dependent increase in the variance of EPSC amplitudes following psilocin wash-in (Levene’s test, P = 0.004; Figure 1E). Gaussian mixture models revealed a bimodal distribution of responses in the 300 nM and 1 μM groups (Figure 1F) and stratification of these data according to the model revealed a group of L5 neurons in which excitatory synaptic input was potentiated (main effect of [psilocin]: P = 0.0002; Figure 1G), and a group in which excitatory synaptic drive was depressed (main effect of [psilocin]: P = 0.0002; Figure 1H), an effect that was more pronounced at higher concentrations (Figure 1I). These data reveal that psilocin’s actions are not uniformly excitatory but bidirectional and cell-type-specific, suggesting a redistribution of synaptic drive within the mPFC microcircuitry[30].

### Cortico-amygdala projections are selectively inhibited by psilocin, dependent on both 5HT_2A_ and 5HT_1A_ receptor subtypes

Given psilocybin’s effects in the ABT, we asked whether these distinct neuronal populations differentially contributed to modulation of affective biases. Retrograde labelling of cells projecting from PrL to the basolateral nucleus of the amygdala (BLA) (Figure 2A) enabled the concurrent recording of EPSCs in L5 cortico-amygdala (CA) and L5 unlabeled control cells.

**Figure 2.**
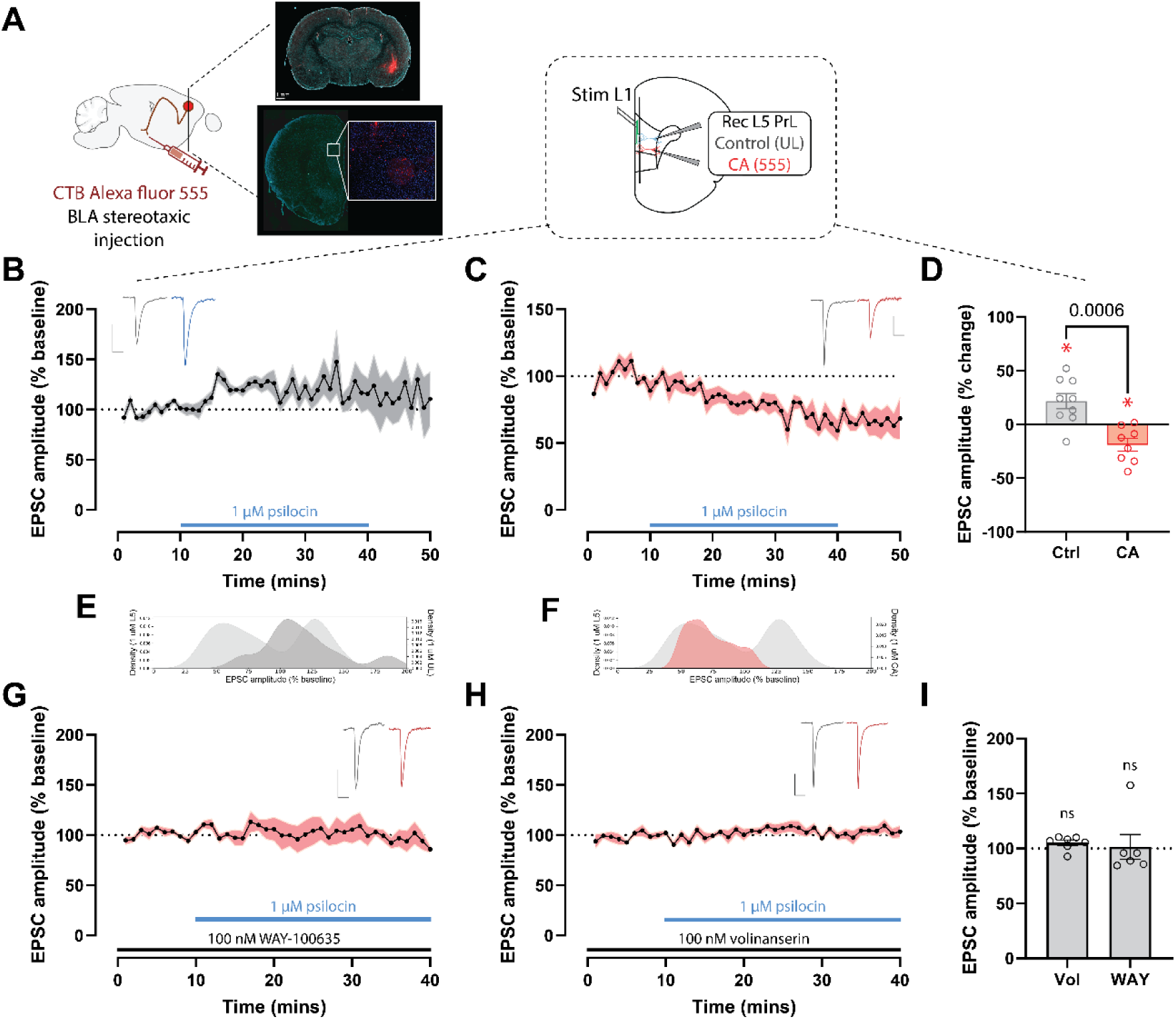
Excitatory input to cortico-amygdala projections are selectively down-regulated by psilocin, depending on 5HT_2A_ and 5HT_1A_ receptors. **A)** Cortico-amygdala (CA) neurons were labelled using the retrograde fluorescent tracer cholera toxin subunit b (CTb) conjugated to Alexa-fluor 555, allowing the recording of CA cells and non-labelled control cells. **B)** Time-series data of excitatory synaptic input in non-labelled control cells (n = 9) during application of psilocin (1 µM). Data are normalized to baseline EPSC amplitude. **C)** Time-series data of excitatory synaptic input in labelled CA cells (n = 8) during application of psilocin (1 µM). Data are normalized to baseline EPSC amplitude. **D)** Amplitude of excitatory post-synaptic currents (EPSCs) measured during the final 10 minutes of wash-in between control and labelled CA cells during psilocin (1 µM) exposure. Ratio paired two-tailed t-test versus baseline (NT: t₈ = 2.975, *P < 0.05, CA: t₇ = 2.975, P < 0.05). Welch’s two-tailed t-test to compare between groups: t₁₄.₇₉ = 4.41 **E)** Distributions of EPSC amplitudes during the final 10 minutes of psilocin wash-in in non-labelled control cells overlayed onto the bimodal distribution from the previous experiment. **F)** Distributions of EPSC amplitudes during the final 10 minutes of psilocin wash-in in labelled cortico-amygdala cells overlayed onto the bimodal distribution from the previous experiment. **G)** Time-series data of EPSC amplitudes measured in labelled CA cells (n = 6) during pre-administration of the selective 5-HT_1A_ receptor antagonist WAY-100635 (100 nM) and subsequent psilocin application (1 µM). **H)** Time-series data of EPSC amplitudes measured in labelled CA cells (n = 7) during pre-administration of the selective 5-HT_2A_ receptor antagonist volinanserin (100 nM) and subsequent psilocin application (1 µM). Representative traces in (C,E,G,H) are averaged across 10 minute baseline and final 10 minutes of psilocin wash-in. Scale bars are x = 50 ms and y = 50 pA. **I)** EPSC amplitudes measured during the final 10 minutes of antagonist and psilocin wash-in. One-sample Wilcoxon test versus baseline: (WAY-100635: P = 0.44, Volinanserin: P = 0.11). Data mean ± SEM.

In non-labelled control cells, psilocin selectively induced a potentiation of synaptic input (vs. baseline, P = 0.018; Figure 2B). Strikingly, in CA cells, psilocin selectively induced a depression of excitatory synaptic input (vs. baseline, P = 0.019; Figure 2C), revealing a cell-type specific modulation of synaptic transmission in PrL by psilocin (Figure 2D). Analysis of the distributions of these projection-defined populations revealed overlap between the non-labelled control population and the excited L5 neuronal population from the previous experiment (Figure 2E). On the other hand, the distributions of the CA population and the inhibited population showed considerable overlap (Figure 2F).

Psilocin, and other psychedelic drugs, have distinct poly-pharmacology, and act on a number of serotonin receptor subtypes at clinical doses. Generally, the therapeutic effects of psilocin have been ascribed to its action at the 5HT2A receptor[31, 32], although the 5HT1A receptor has also been recognized for its importance in emotional regulation[33]. Using subtype-selective antagonists, we aimed to elucidate the receptor specific contributions to the synaptic depression on CA cells induced by psilocin. Pre-administration of the 5HT1A receptor antagonist WAY-100635 (100 nM) blocked the synaptic depression induced by psilocin (Figure 2G). Somewhat surprisingly, pre-administration of the 5HT2A receptor antagonist volinanserin (100 nM) also blocked the downregulation of synaptic input to CA cells (Figure 2H). The convergence of both receptor pathways suggests parallel 5-HT_2A_ and 5-HT_1A_ signaling in mediating projection-specific modulation.

### Psilocybin’s modulation of affective biases depends on both 5HT_2A_ and 5HT_1A_ receptor subtypes

We next sought to establish how 5HT receptor sub-types may contribute to the modulation of affective biases by psilocybin. We have previously shown that systemic psilocybin reverses the valence of negative affective biases (Figure 3A) when measured 24 hours following administration of a RAAD and proposed this is one of the critical features relevant to its sustained antidepressant effects[4]. Similar to the inhibition of CA excitatory input, the behavioral effects of psilocybin (0.3 mg/kg IP and 1 mg/kg IP) at 24 hours were fully blocked by pretreatment with WAY-100635 or volinanserin (Figures 3B-D).

**Figure 3.**
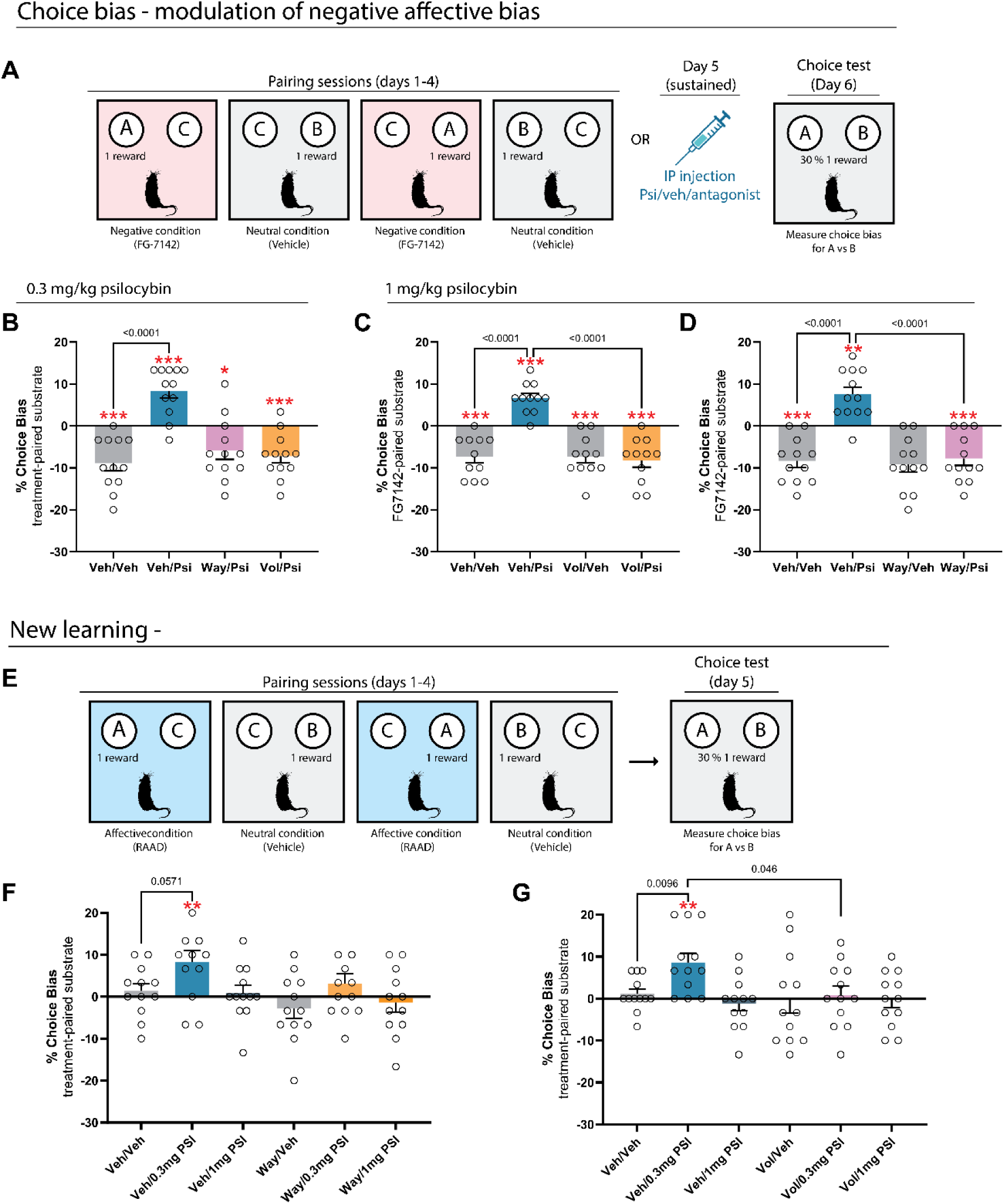
Modulation of affective biases by psilocybin depends on both 5HT_1A_ and 5HT_2A_ receptor subtypes. **A)** Schematic of the affective bias test used to assess sustained effects of psilocybin on a negative affective bias induced by FG-7142 (3 mg/kg SC). Animals were tested 24 h after receiving vehicle/vehicle, vehicle/psilocybin (0.3 or 1 mg/kg IP), or antagonist/psilocybin combinations. **B)** Effects of psilocybin (0.3 mg/kg IP), WAY-100635 (0.1 mg/kg IP), and volinanserin (0.1 mg/kg IP) on retrieval of negatively biased memories. One-sample t-tests vs 0% bias (t₁₁ = 4.93, 5.00, 2.73, 4.57; *P < 0.05, ***P < 0.001) and repeated-measures ANOVA (main effect: F₃,₃₃ = 19.65, P < 0.0001) with S í da k’s multiple comparisons test (n = 12). **C)** Effects of psilocybin (1 mg/kg IP) and WAY-100635 (0.1 mg/kg IP) on retrieval of negatively biased memories. One-sample t-tests vs 0% bias (t₁₁ = 5.33, 4.42, 4.98, 4.69; **P < 0.01, ***P < 0.001) and repeated-measures ANOVA (main effect: F_2, 22_ = 25.96, P < 0.0001) with S í da k’s test (n = 12). **D)** Effects of psilocybin (1 mg/kg IP) with volinanserin (0.1 mg/kg IP). One-sample t-tests vs 0% bias (t₁₀ = 4.92, 6.06, 4.92, 4.98; ***P < 0.001) and repeated-measures ANOVA (main effect: F_3, 30_ = 22.20, P < 0.0001) with S í da k’s test (n = 11). **E)** Schematic of the affective bias test used to assess psilocybin effects on new learning. Animals received vehicle/vehicle, vehicle/psilocybin (0.3 or 1 mg/kg IP), antagonist/vehicle, or antagonist/psilocybin (0.3 or 1 mg/kg IP) before each independent substrate–reward association learning session. **F)** Effects of the 5-HT_1A_ antagonist WAY-100635 (0.1 mg/kg IP) on psilocybin-induced bias during new learning. Choice bias was quantified 24 h after the final pairing session. Data were analyzed with a one-sample t-test vs 0% bias (vehicle/psilocybin 0.3 mg/kg IP: t₁₂ = 3.12, **P < 0.01) and a repeated-measures ANOVA (main effect of treatment: F_4, 43_ = 2.63, P = 0.484), followed by S í da k’s multiple comparisons test (n = 12). **G)** Effects of the 5-HT2A antagonist volinanserin (0.1 mg/kg IP) on psilocybin-induced bias in new learning. One-sample t-test vs 0% bias (vehicle/psilocybin 0.3 mg/kg IP: t₁₁ = 3.87, **P < 0.01) and repeated-measures ANOVA (main effect of treatment: F_5, 55_ = 2.67, P = 0.0313) with S í da k’s multiple comparisons test (n = 12). Data are mean ± SEM.

We have also previously shown that psilocybin and conventional SSRI antidepressants share the ability to positively bias new experiences[4, 16, 18] so we also tested whether these effects were modulated by 5HT_2A_ and 5HT_1A_ receptors. By administering psilocybin before one of the substrate-reward pairing sessions (Figure 3E) we confirmed previous findings that low-dose psilocybin positively biases new reward memories, similar to conventional antidepressants. Strikingly, here we find that either WAY-100635 (Figure 3F) or volinanserin (Figure 3G) administered systemically before psilocybin are sufficient to block the positive bias induced by psilocybin in the ABT.

### Prelimbic psilocin infusion causes sustained modulation of negative affective biases, parallel to increased inhibition to CA projecting neurons

Infusion of psilocin directly into PrL was sufficient to effectively reverse a negative affective bias induced by FG-7142 24 hours after infusion (Figures 4A and 4B), an effect that recapitulates the positive affective modulation and re-learning effect previously seen with systemic psilocybin[4]. Therefore, PrL is a critical region in mediating the sustained modulation of affective biases by psilocybin, and plasticity of CA cells may underlie this.

**Figure 4.**
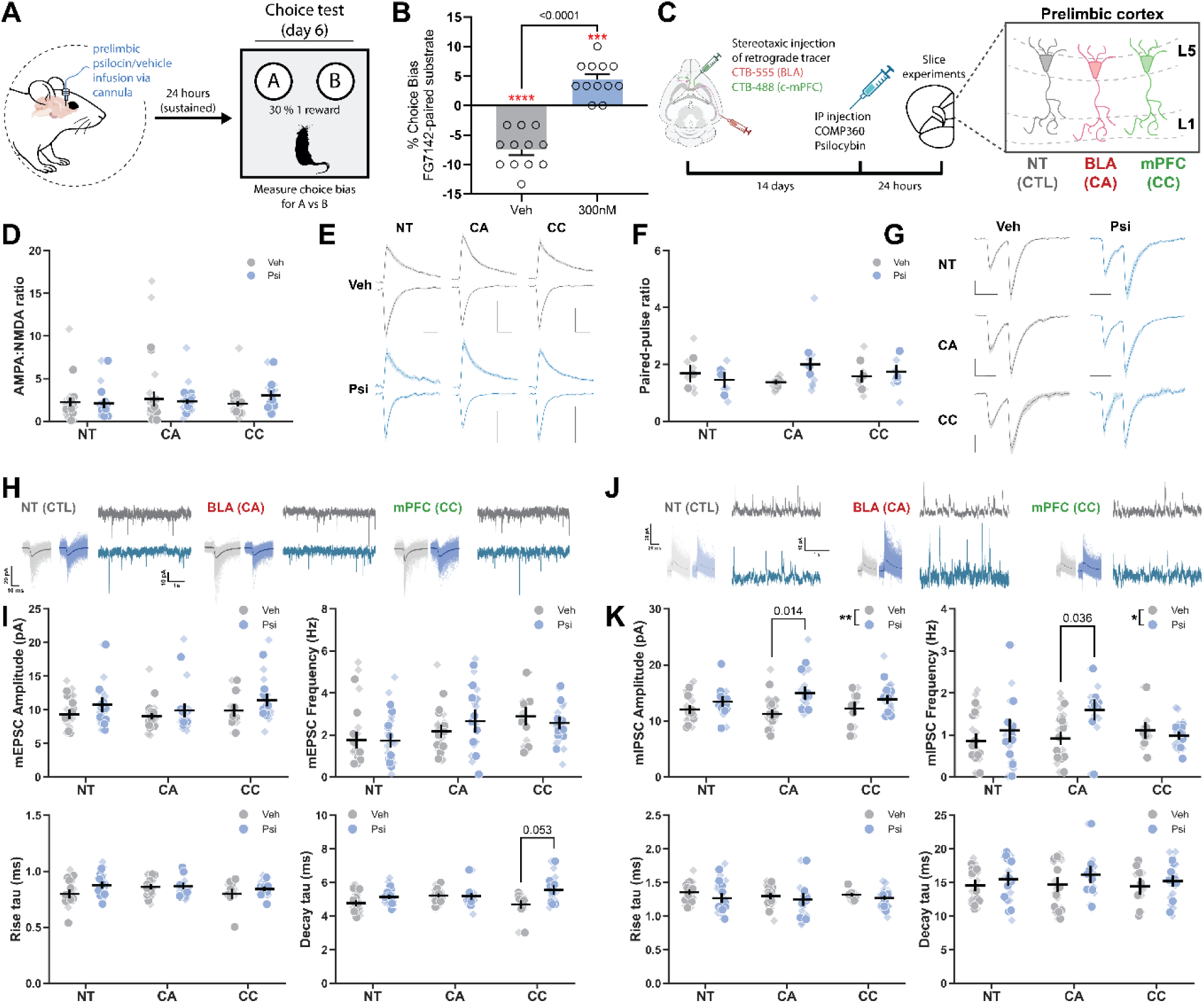
Psilocin infusion replicated the systemic behavioral effects of psilocybin in the ABT 24 hours post-treatment, parallel to sustained inhibition of cortico-amygdala projections. **A)** Overview of the affective bias test (ABT) protocol used to assess modulation of a negatively biased memory 24 hours after prelimbic infusion of vehicle or psilocin (300 nM). **B)** Choice biases in male rats (n = 12 per group) 24 hours after prelimbic infusion of vehicle or 300 nM psilocin. Data were analyzed using one-sample t-tests against a null hypothesis mean of 0% choice bias (Veh: t₁₁ = 8.074, ****P < 0.0001, Psilocin: t_11_ = 5.204, ***P < 0.001) and a paired t-test to test between conditions (t_11_ = 11.46, P < 0.0001). **C)** Labelling strategy using CTb–Alexa Fluor 555 and CTb–Alexa Fluor 488 to identify cortico-amygdala (CA), cortico-cortical (CC), and non-tagged (NT) neurons for electrophysiological recordings. Whole-cell recordings were obtained 24 hours after systemic administration of 0.3 mg/kg IP psilocybin or vehicle. **D)** AMPA:NMDA ratios were calculated from excitatory post-synaptic currents (EPSCs) recorded by voltage clamping cells at -70 and +40 mV. A linear mixed model (LMM) was used to test for drug, projection, and interaction effects (n = 16 (NT), 19 (CA), 12 (CC) cells from 12 vehicle treated animals, n = 14 (NT), 16 (CA), 11 (CC) from 12 psilocybin treated animals). **E)** Representative EPSC traces averaged across experiments. Scale bar x = 50 ms and y = 100 pA. **F)** Paired-pulse ratios were calculated from EPSCs stimulated at an interval of 50 ms recorded at -70 mV. LMM used to test for drug, projection, and interaction effects (n = 6 (NT), 8 (CA), 7 (CC) cells from 6 vehicle treated animals, n = 5 (NT), 9 (CA), 6 (CC) from 6 psilocybin treated animals). **G)** Representative EPSC traces averaged across experiments. Scale bar x = 50 ms and y = 100 pA. **H)** Representative traces and average waveforms of miniature EPSC (mEPSC) recordings and events, respectively, from CA, CC, and NT neurons. **I)** The amplitude, frequency, and kinetics of mEPSCs were analyzed 24 hours after treatment with vehicle or 0.3 mg/kg IP psilocybin. LMM was used to test for treatment, projection, and interaction effects (decay tau: effect of Treatment X Projection interaction; P = 0.012) (n = 16 (NT), 18 (CA), 9 (CC) cells from 12 vehicle treated animals, n = 16 (NT), 16 (CA), 15 (CC) from 12 psilocybin treated animals). **J)** Representative traces and average waveforms of miniature IPSC (mIPSC) recordings and events, respectively, from CA, CC, and NT neurons. **K)** The amplitude, frequency, and kinetics of mIPSCs were analyzed 24 hours after treatment with vehicle or 0.3 mg/kg IP psilocybin. LMM was used to test for treatment, projection, and interaction effects (amplitude: main effect of drug; P = 0.005). Simple comparisons were made post-hoc and significance corrected for multiple comparisons using the Bonferroni method (n = 16 (NT), 19 (CA), 9 (CC) cells from 12 vehicle treated animals, n = 14 (NT), 13 (CA), 16 (CC) from 12 psilocybin treated animals). Data are presented as superplots with individual cells shown as semi-transparent diamonds and animal means shown as solid circles. The plotted lines represent mean ± SEM of animal averages.

To test this, we again utilized a retrograde labelling strategy, expressing CTb-555 in CA cells and CTb-488 in neighboring cortico-cortical (CC) cells, enabling the recording in ex-vivo preparations of CA, CC, and non-tagged (NT) control cells, 24 hours after a single low-dose of psilocybin (0.3 mg/kg IP) or vehicle (Figure 4C).

Contrary to the acute effects of psilocin, we found no evidence of long-term changes to evoked excitatory currents (Figure 4D-E) nor any changes to pre-synaptic glutamate release properties measured by paired-pulse ratios (Figure 4F-G). Intrinsic membrane and action potential firing properties were also unchanged in all cell types (Figure S2).

In line with the unchanged evoked EPSC properties, miniature EPSC analyses (Figure 4H) revealed no changes to the frequency or amplitude of mEPSCs in any cell type. However, we observed an interaction between psilocybin treatment and cell-type on mEPSC decay kinetics (P = 0.012, LMM), which was driven by a selective increase in the decay tau in CC neurons (Figure 4I). These data indicate that psilocybin modulates excitatory synaptic kinetics in a circuit-specific manner, selectively targeting corticocortical rather than cortico-amygdala pathways.

Strikingly, analysis of miniature inhibitory post-synaptic current (mIPSC) properties (Figure 4J) revealed a significant main effect of psilocybin on mIPSC amplitude (P = 0.005, LMM) and frequency (P = 0.037, LMM). Post-hoc comparisons confirmed that this increase in inhibitory drive, observed in both amplitude and frequency, was restricted to the CA population (Figure 4K). This enhanced inhibition of CA projections induced by psilocybin could underlie a longer-term suppression of excitatory output in the CA circuit, contributing to the reduced activity in this network and the sustained modulation of affective biases observed in the ABT.

### Chemogenetic inhibition of CA cells replicates psilocybin’s modulation of affective biases

To directly test if enhanced inhibition of the CA circuit modulates affective biases, we used a dual viral approach to selectively express a Gi coupled inhibitory DREADD in the CA projection neurons (Figure 5A).

**Figure 5.**
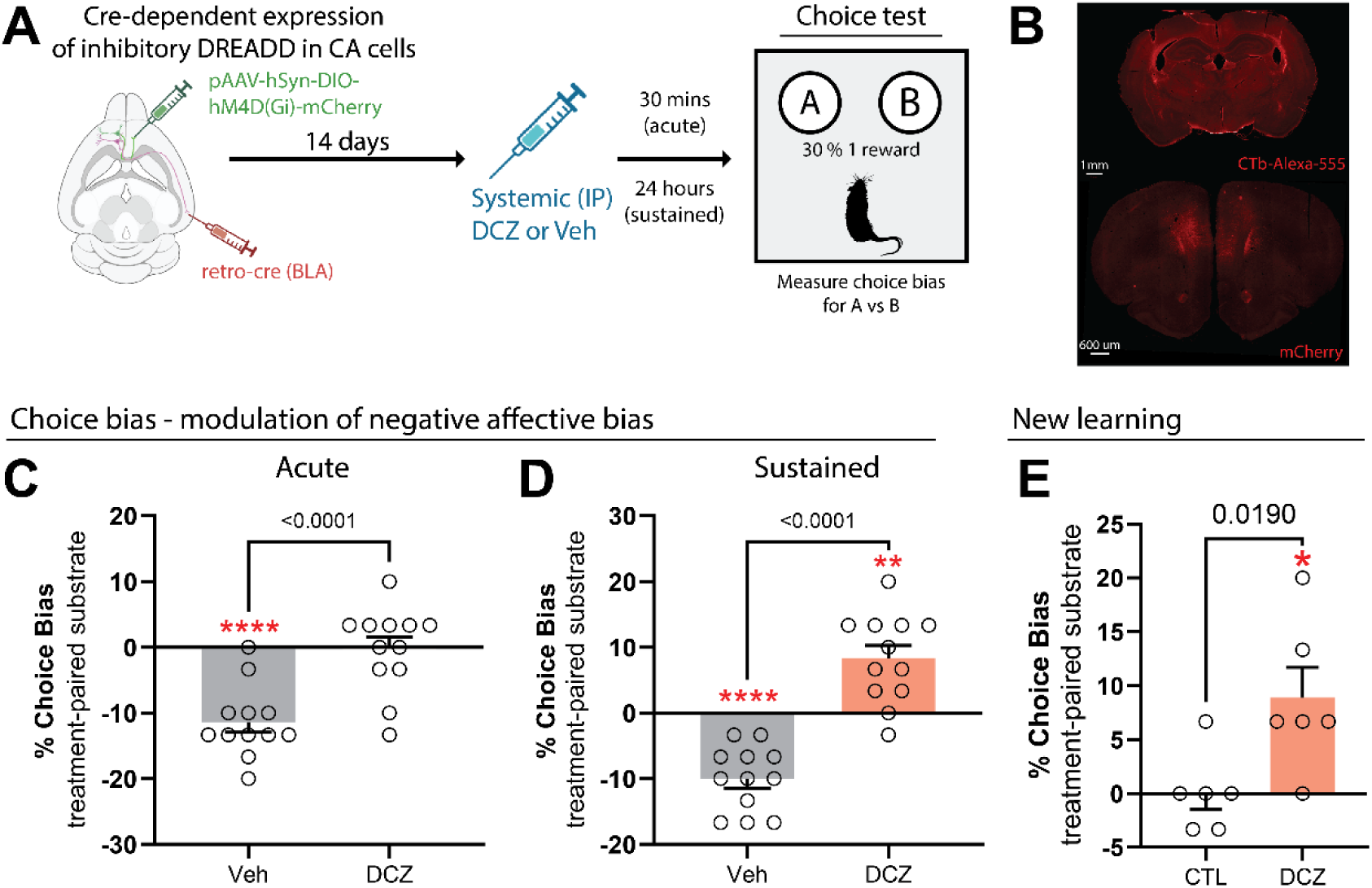
Chemogenetic inhibition of CA cells replicates the modulation of affective biases induced by low-dose psilocybin. **A)** Inhibitory Gi-coupled DREADDs were expressed selectively in CA cells by the injection of retro-cre and pAAV-hSyn-DIO-hM4D(Gi)-mCherry viruses to BLA and PrL, respectively. **B)** Representative image of CTb-AlexaFluor-555 expression in the BLA and mCherry in prelimbic medial prefrontal cortex. **C)** Acute modulation of a negative affective bias induced by FG-7142 (3 mg/kg SC) was measured following oral administration (PO) with 0.1 mg/kg descoclozapine (DCZ) or vehicle. Choice bias was quantified 1 h after the final pairing session. Data were analyzed with a one-sample t-test vs 0% bias (vehicle: t_11_ = 7.3, ****P < 0.0001) and a paired t-test between groups (t_11_ = 6.701) (n = 12). **D)** Sustained modulation of a negative affective bias was tested 24 hours after administration of vehicle or DCZ. Data were analyzed with a one-sample t-test vs 0% bias (vehicle: t_11_ = 7.036, ****P < 0.0001, DCZ: t_11_ = 4.282, **P < 0.01) and a paired t-test between groups (t_11_ = 8.848) (n = 12). **E)** The formation of an affective bias in the learning of a new reward memory by intraperitoneal (IP) injection of DCZ during the pairing sessions was also tested as described above. New learning was assessed either before viral expression of DREADDs in CA cells (CTL), or following viral expression of DREADDs in CA cells (DCZ). Data were analyzed using a one-sample t-test vs. 0% choice bias (t_5_ = 3.162, *P < 0.05) and an unpaired t-test between groups (t_10_ = 2.794)(n=6). Data are mean ± SEM.

Consistent with our predictions, inhibition of this circuit, in a manner analogous to the actions of psilocybin demonstrated in the electrophysiological analyses, replicated the acute attenuation of a negative affective bias (One sample t-test vs. 0: Veh P<0.0001, DCZ ns; Figure 5C). Administration of the DREADD agonist descoclozapine (DCZ) to inhibit the CA neurons was also sufficient to facilitate the re-learning of the negative affective bias 24 hours after administration (One sample t-test vs. 0: Veh P < 0.0001, DCZ P = 0.0013; Figure 5D) and positively bias the learning of new reward memories (One sample t-test vs. 0: CTL ns, DCZ P = 0.025; Figure 5E), effectively fully recapitulating the effects of psilocybin on the modulation of affective biases.

## Discussion

The present findings reveal a novel, circuit-specific mechanism by which a single dose of the serotonergic psychedelic psilocybin modulates affective biases, namely, the suppression of excitatory drive to cortico-amygdala (CA) neurons in prelimbic mPFC.

We show that psilocin infusion into the PrL is sufficient to reproduce the acute and 24 hour post-treatment effects of systemic psilocybin on negative affective biases[4], identifying this mPFC subregion as a central locus of the behavioral effects in the ABT. Psilocin acutely suppresses excitatory drive onto CA neurons through both 5-HT_2A_ and 5-HT_1A_ receptor signaling, and both receptor subtypes are also required for modulation of affective biases observed in the ABT. Chemogenetic inhibition of CA neurons with G_i_-coupled DREADDs replicated psilocybin’s behavioral effects in the ABT, acutely attenuating negative affective bias, facilitating re-learning of a negatively biased memory with a more positive affective valence, and positively biasing new reward memories. These findings demonstrate that suppression of activity in PrL–BLA projections is sufficient to mediate both the rapid and enduring components of psilocybin’s behavioral effects and establish CA neuron inhibition as a key circuit substrate of these therapeutic effects.

Our data provide a mechanistic explanation for how psilocybin produces rapid and sustained changes to affective biases, a critical neuropsychological mechanism implicated in MDD and antidepressant efficacy[34]. Prefrontal connectivity with the amygdala is central in the control of emotional behavior and memory[35–38] and optogenetic inhibition of the dorsal medial PFC to BLA projection was recently found to decrease anxiety like behaviors and induce effects predictive of antidepressant efficacy[39]. Our results extend these findings by demonstrating that inhibition of CA neurons with targeted psilocin infusion, and chemogenetic inhibition, is sufficient to acutely attenuate previously learned negatively biased memories. In addition, enhanced suppression of CA cells observed at 24 hours post-treatment in the electrophysiology experiments suggests a mechanism for longer-term modulation of affective biases, potentially longer-term adaptation in CA circuits enabling more positive affective processing. These behavioral dynamics are parallel to observations in humans that psilocybin produces shifts towards more positive emotional interpretation[40], enhanced emotional openness and social recognition, and improved cognitive flexibility[41], all of which are associated with therapeutic outcomes.

At the systems level, the suppression of PrL-BLA projections provides a plausible neural substrate for the shifting of network activity away from maladaptive, persistently-engaged negative affect networks[42, 43], and towards a more flexible cortico-limbic configuration. Human neuroimaging studies similarly highlight the importance of prefrontal-amygdala connectivity in affective processing and report psilocybin-induced modulation of functional coupling between these regions[29, 44]. While recent findings suggest that such large-scale network changes may emerge from cell- and projection-specific mechanisms within mPFC microcircuits [20, 30], linking these levels of analysis remains a challenge. Blood oxygenation level dependent (BOLD) functional MRI provides critical systems-level insight, though the spatial and temporal resolution afforded by this approach may not fully capture the subregion and even subcellular differences in information processing that serotonergic psychedelics introduce. Furthermore, recent work demonstrating a dissociation between hemodynamic and neuronal responses following serotonin receptor agonism[45, 46] emphasizes the importance of multimodal approaches that integrate high-resolution circuit analysis with larger scale systems analyses to bridge the cellular and network level changes induced by psychedelic drugs.

At the cellular level, psychedelic drugs, including psilocybin, have been shown to promote the rapid and sustained growth of dendritic spines and branches [19, 27], effects commonly attributed to activation of 5HT_2A_ receptors[47], although the precise cellular and molecular mechanisms remain incompletely understood[48, 49]. Our findings suggest that antidepressant-like actions of psilocybin may not arise from global cortical excitation or indiscriminate structural plasticity, but instead from a rebalancing of prefrontal-limbic circuit dynamics. This interpretation aligns with recent reports of activity-dependent input reorganization in the anterior cingulate cortex (ACC) following psilocybin exposure in the mouse[30], in which connections to pyramidal-tract (PT) and intra-telencephalic (IT) neurons were differentially modified in an experience-dependent manner[20, 30]. While our data focus on distinct prelimbic projection neurons, the principle of selective, projection-specific circuit remodeling as a substrate for behavioral change is consistent across these studies. Notably, even within mPFC, distinct subregions and their projections exert opposing influences on affective behavior, PrL projections to the BLA promote fear expression, whereas projections to the BLA from the neighboring infralimbic cortex (IL) facilitate fear extinction and suppression[50–52].

Given the proximity and connectivity between PrL and ACC, it is plausible that partially overlapping, but functionally distinct, PT and IT populations contribute to affective regulation through complementary pathways. Indeed, both *Fezf2*-(IT) and *Plxnd1-*expressing neurons in PrL send projections to the BLA[53], suggesting that classification by projection target may stray from transcriptional definitions of PT and IT cells. Determining whether CA neurons that are suppressed by psilocybin treatment represent a transcriptionally distinct subtype that could confer even greater therapeutic potential[54] will be an important avenue for future work. Similarly, while electrical stimulation of Layer I effectively recruits broad excitatory afferent drive to these cells, it inherently activates a heterogeneous population of fibers, including inputs from the mediodorsal thalamus, contralateral mPFC, and BLA[55]. Dissecting exactly which of these input pathways are selectively depressed by psilocin, and identifying the specific local interneuron populations mediating the enhanced inhibition observed at 24 hours, represent critical next steps to fully resolve the synaptic architecture of this therapeutic mechanism.

The receptor mechanisms underlying these effects appear similarly nuanced, contrary to the theory that the 5HT_2A_ receptor is the predominant mediator of psilocybin’s therapeutic effects[20, 21, 31, 47, 56]. The present study indicates that coordinated 5HT_2A_ and 5HT_1A_ signaling within defined microcircuits is critical, rather than a single excitatory pathway promoting global plasticity effects. Indeed, blocking either receptor subtype prevented both synaptic effects in CA cells and behavioral effects in the ABT. Cortical (post-synaptic) 5HT_1A_ receptors appear to play a critical role in positive effects on stress-coping behaviors[57], and recently specific pharmacological targeting of these heteroreceptors has been shown to produce similar positive behavioral effects[58]. Therefore, it seems that the unique poly-pharmacology of psychedelic drugs like psilocybin could confer them with the ability to rebalance cortical microcircuits via combined action at many receptor subtypes[23], but critical targets may be 5HT_2A_ and 5HT_1A_ receptors.

Together, these data suggest that psychedelics such as psilocybin may act therapeutically not by globally amplifying neuroplasticity, but by reconfiguring maladaptive emotional networks through selective serotonergic modulation of prefrontal-limbic circuits. Such cell- and circuit-specific actions may represent a unifying mechanism underlying rapid-acting antidepressant efficacy, and defining how these maladaptive networks manifest and are remodeled in depression could reveal new targets for precision therapies in mood disorders.

## Methods

### Animals and housing

Male Lister Hooded rats (Envigo, UK) were housed in pairs in enriched laboratory cages (55 × 35 × 21 cm) containing aspen woodchip bedding, paper bedding, a cotton rope, wood block, cardboard tube (Ø 8 cm) and a red Perspex house (30 × 17 × 10 cm). Housing was maintained under temperature-controlled conditions (21 ± 1 °C) on a 12:12 h reverse light–dark cycle (lights off at 08:00 h). Rats had ad libitum access to water and were maintained at ∼90% of their free-feeding weight by daily rationing of laboratory chow (∼18 g/per rat, Purina, UK) provided in the home cage at the end of each experimental day. Daily health and welfare checks were carried out by animal facility staff and researchers. Behavioural work was conducted during the active phase (09:00–17:00 h). All procedures complied with the UK Animals (Scientific Procedures) Act 1986 and were approved by the University of Bristol Animal Welfare and Ethical Review Body and the UK Home Office (PPL PP9516065).

Rats from five independent cohorts (n = 12 per cohort; details in Supplementary Table S1) were used in the behavioural experiments. At the start of training, rats were 10–11 weeks old, weighing between 300–350 g. Sample size was informed by our previous affective bias test studies and a meta-analysis[4, 17, 59] demonstrating similar affective biases. Given that similar effect sizes were observed regardless of strain or sex, we used only males to reduce total animal numbers.

### Affective Bias Test: training and testing

Before testing, rats underwent pre-training handling (https://www.3hs-initiative.co.uk/) and arena habituation following the previously published protocol[60]. In the second week, over five consecutive days, animals learned to dig in ceramic bowls filled with sawdust, with task difficulty gradually increasing. Training concluded with a novel substrate discrimination session, where rats had to choose and dig in the substrate associated with a food reward. ‘Correct’ choices were recorded when the reward-paired substrate was selected, ‘incorrect’ choices when the unrewarded substrate was chosen, and ‘omissions’ when the rat failed to approach within 10s. The criterion for completion was six consecutive correct trials within 20 total trials. Once successfully trained, animals undergo a reward learning assay[60] to confirm that they would develop a reward-induced positive bias.

Testing weeks, each consisted of four consecutive pairing sessions generating two distinct substrate–reward associations (one under control conditions and one under treatment) (for details, see Figure 1). On the fifth day (acute modulation of negative affective biases, the new learning studies or reward learning assay) or sixth day (sustained modulation of negative affective biases), a choice test assessed memory retrieval with or without drug pre-treatment (see Figure 1). During the choice test, both previously treatment-paired rewarded substrates were presented simultaneously over 30 randomly reinforced trials. Latency to make a choice were recorded. Each drug was tested as an independent experiment and a within-subject design was used, with the experimenter blind to treatment and with a fully counterbalanced experimental design.

### Modulation of the affective biases by the antagonist compounds

These experiments used a within-subject design and were carried out over of four weeks in the sustained modulation studies, while the new learning studies over 6 weeks. To induce a negative bias, all animals received the same procedure, on treatment days, they were injected with FG7142 or vehicle 30 min. prior to substrate-reward pairing session (see Fig. 3). In the new learning experiments, all animals (cohort 1) received two injections: first with either vehicle, volinanserin 0.1mg/kg IP, or WAY100635 0.1mg/kg IP, and 15 minutes later with either vehicle or psilocybin. These injections were administered 60 minutes before the pairing sessions, followed by a choice test on day 5. To test the effects of antagonists on the sustained modulation of biases, rats from cohort 2 were first treated with FG7142 or vehicle prior to pairing sessions to induce a negative bias. On day 5, each animal received two injections (vehicle, volinanserin 0.1mg/kg IP, or WAY100635 0.1mg/kg IP), followed 15 minutes later by a second injection of either vehicle or psilocybin (0.3 or 1.0mg/kg IP). The choice test was conducted on day 6, 24 hours after the final injection.

### Medial prefrontal cortex cannulation and infusion studies

For experiments involving infusion of psilocin into the prelimbic cortex, male rats from cohorts 3 were first implanted with a bilateral guide cannula (32-gauge, Plastics One, UK) into the medial prefrontal cortex (stereotactic coordinates from Bregma [+2.70mm anterior/posterior (AP), ±0.80mm medial/lateral (ML) and −2.1mm dorsal/ventral (DV) from dura]. For surgeries rats were anaesthetized with 2–5% isoflurane first in a chamber, then mounted on a stereotaxic apparatus with a heating pad set at 37 °C below the rat. The skull was levelled, and two craniotomies were performed. Guide cannulae were implanted and secured to the skull by 3 stainless steel screws and a head mount built with dental cement (dePuy, Johnson&Johnson, UK). The guide cannula was protected by a dummy cannula and a dust cap to prevent any infections. After the two weeks recovery period, all animals were habituated to the infusion procedure during two sessions on separate days. During experimental infusions, each rat was lightly restrained (see https://www.3hs-initiative.co.uk/) while the dummy cannula was removed, the injector was placed through the guide cannula for a 1 minute pre-infusion, 2 minutes for infusion of vehicle (PBS) or drug (300 nM, 1 uM psilocin, 1.0 μl per site, with a flow rate of 0.5 μl per minute) and for 2 minutes post-infusion to allow diffusion of the vehicle/drug into the surrounding tissue. All animals were infused with either vehicle or psilocin in a fully counterbalanced design. Animals were then tested, either immediately post treatment for the acute modulation of a negative affective bias or 24 hours post treatment for sustained modulation of a negative affective bias. At the end of the study, cannulated rats were killed by transcardiac perfusion with PBS followed by 4% paraformaldehyde under terminal sodium pentobarbitone anaesthesia and the brain was removed, sectioned and stained with cresyl violet to determine cannula position. All animals were included in the post-histological verification analysis.

### Drugs

The compound used to induce a negative affective bias in rats was FG7142, benzodiazepine inverse agonist (3 mg/kg, SC). FG7142 induces a reliable negative affective bias in the ABT and has rapid washout limiting the potential for adaptive or cumulative effects which is beneficial for within-subject study designs run over multiple weeks. The compounds tested were COMP360 (COMP360 – Compass Pathways proprietary synthetic form of crystalline psilocybin) (0.3, and 1.0 mg/kg; IP; t= -60min and -24 h), volinanserin (0.1 mg/kg; IP; t= -75 min and -24h), WAY100635 (0.1 mg/kg; IP; t= -75min and -24h), DCZ (0.1 mg/kg, PO, t=-60 min and -24h) and psilocin (300 nm, 1 μM; 1.0 μl per site, for more details see Table S1). The doses used were based on previous publications and estimates of receptor occupancy where available. COMP360 psilocybin was supplied by COMPASS Pathways, WAY100635 was purchased from HelloBio, and volinanserin and psilocin from Cambridge Bioscience. FG7142 was suspended in 0.9% saline with a drop of Tween 80 added to maintain suspension; psilocybin and WAY100635 was dissolved in 0.9% saline and pH adjusted; volinanserin was dissolved in 5% DMSO, 10% Cremophor and 85% saline. DCZ was dissolved in 10% condensed milk. All drugs were freshly prepared every day, and they were administered in a dose volume of 1.0 ml/kg. For FG7142-induced negative affective biases we selected doses previously reliable in the affective bias test[17, 60]. The doses for psilocybin were based on previous studies[60]. Intraperitoneal and subcutaneous injections were done by using a low-stress, non-restrained methods developed in our research group (https://www.3hs-initiative.co.uk/). Prior to the start of the study, all rats were habituated to the holding positions required for IP and SC dosing. In studies testing psilocybin, the number of head twitches and wet dog shakes were scored (see Table S1).

### Brain slice preparation

Rats were terminated by overdose of isoflurane and death confirmed by cervical dislocation, and the brain rapidly removed and placed into ice-cold sucrose modified artificial cerebrospinal fluid (aCSF) containing (in mM): 52.5 NaCl, 2.5 KCl, 25 NaHCO3, 1.25 NaH2PO4, 5 MgCl2, 25 D-Glucose, 100 sucrose, 2 CaCl2, 0.1 kynurenic acid, saturated with 95% O2/5% CO2. The brain was blocked and slices (300 μm) cut at a modified coronal axis using a vibrating blade microtome (Leica VT1000 S). Slices were placed in a holding chamber containing aCSF containing (in mM): 124 NaCl, 3 KCl, 26 NaHCO3, 1.4 NaH2PO4, 1 MgSO4, 10 D-Glucose, 2 CaCl2, where they incubated at 32 °C for 30 minutes, and then at room temperature for at least 30 minutes, before being transferred to the recording chamber.

### Whole-cell Electrophysiology

Brain slices were visualized using DIC optics and whole-cell recordings were taken from pyramidal neurons in L5 of the prelimbic cortex. For studies of labelled cells, cells were first identified using fluorescence microscopy and subsequently patched using DIC. Patch electrodes with a resistance of 3–6 MΩ were pulled from borosilicate filamented glass capillaries (1.5 OD × 0.86 ID × 100 L mm, Harvard Apparatus) with a horizontal puller (P-97, Sutter Instrument Co., UK) and filled with internal solution. For the recording of synaptic events the internal solution contained (in mM): 8 NaCl, 130 CsMeSO4, 10 HEPES, 0.5 EGTA, 4 MgATP, 0.3 NaGTP, and 5 QX-314. For the recording of intrinsic membrane and action potential properties the internal solution contained (in mM): 135 K-Gluconate, 8 NaCl, 10 HEPES, 2 MgATP, and 0.3 NaGTP. AMPA and NMDA receptor mediated currents (for AMPA:NMDA ratios) were recorded in the presence of 100 μM picrotoxin (HelloBio, UK) to block GABA_A_ currents, and while the cell was held at -70 and +40 mV, respectively, in voltage-clamp configuration. Miniature post-synaptic currents were recorded in the presence of 1 μM tetrodotoxin citrate (HelloBio, UK) to block action-potential mediated events. mEPSCs were recorded and -70 mV and mIPSCs were recorded at 0 mV. For current-clamp recordings of membrane and spiking properties, a small current (< 100 pA) was injected such that membrane potential was - 70 mV, and from here a series of hyperpolarising and depolarising currents were injected via the recording electrode. Whole cell recordings were made with a MultiClamp 700B amplifier (Molecular Devices, USA), filtered at 2.4 kHz and digitised at 10 kHz (VC) or unfiltered and digitized at 40 KHz (IC) with a BNC-2110 interface (NI, USA) and WinLTP acquisition software[61]. Series resistance was monitored throughout all experiments and cells that showed >40% change were discarded from subsequent analysis. Recordings were also rejected from analysis if the series resistance was greater than 30 MΩ. Analyses were performed using WinLTP software, Clampfit 9.2 (Molecular Devices, USA), and custom Python code.

### Stereotaxic surgeries for retrograde labelling and viral injections

For chemogenetic and retrogtade labelling surgeries, rats (cohort 4 and 5) were anesthetized with 2–5% isoflurane, initially in an induction chamber and then maintained under anesthesia while mounted on a stereotaxic apparatus. A heating pad set to 37 °C was placed beneath the animals to maintain body temperature. After leveling the skull, four craniotomies were performed. Stereotaxic coordinates for dual-site injections, relative to Bregma, were as follows: for the medial prefrontal cortex (mPFC), +2.60/+3.20 mm anterior-posterior (AP), ±0.60 mm medial-lateral (ML), and - 4.00 mm dorsal-ventral (DV) from the dura; for the basolateral amygdala (BLA), -2.20/- 3.00 mm AP, ±4.60 mm ML, and -8.60 mm DV.

For retrograde labelling studies, unilateral, dual-site infusions into the BLA were performed using 300 nL of cholera-toxin-subunit-B (CTB) conjugated to Alexa Fluor 555 and unilateral, single-site infusions of CTB-Alexa Fluor 448 (Invitrogen, USA) into contralateral (relative to recording) prelimbic mPFC. CTB was infused at a rate of 50 nL/s.

For DREADDs experiments, bilateral, dual-site infusions into the BLA were performed using 300 nL of a Cre-promoter virus (AAVrg-EF1a-Cre, Addgene #55636, titer: 7 x 10^12^ vg/ml). The first 50 nL of virus was delivered at a speed of 500 nL/s, with the remaining volume infused at 50 nL/min. Subsequently, 300 nL of mCherry-labeled DREADDs virus (AAV5-hSyn-DIO-hM4D(Gi)-mCherry, Addgene #44362, titer: 7 x 10^12^ vg/ml) was injected bilaterally into the mPFC at the same infusion rates as used for the BLA. All infusions were made using an infusion pump (World Precision Instruments, USA) and the Hamilton microinjection syringe left in place for 5 minutes post-infusion to allow for diffusion and to minimize backflow.

Rats were allowed to recover for two weeks post-surgery to ensure adequate viral/tracer expression before proceeding with behavioral/electrophysiology experiments. At the end of the DREADDs study, all animals similarly to cohort 3 were euthanised and post-histological verification of the viral expression was analysed. For the electrophysiology experiments, the part of the brain containing the BLA was placed in 4% PFA immediately after brain extraction, and histological confirmation of the injection site later performed. Confirmation of contralateral mPFC injection sites was performed during recording sessions.

### Data and statistical analysis

Data were analysed and figures were created using GraphPad Prism 10.2.0 (GraphPad Software, USA) and custom Python code (Pandas, Scipy, Statsmodels). Data in the text and in the figures are presented as mean ± SEM unless otherwise stated. The level of significance was assigned * if p < 0.05, ** if p < 0.01, *** if p < 0.001 and **** if p<0.0001 for statistical comparisons of all datasets. Sample sizes for behavioral experiments were guided by anticipated medium-to-large effect sizes for drug- and reward-induced biases in Lister Hooded rats, as established in previous studies[16, 18, 59] and a recent meta-analysis[17]. For ex-vivo slice experiments, sample sizes were determined based on established empirical standards in previously published studies using similar experimental setups[62, 63].

### Behavioural data analysis

Choice bias score was calculated as the number of choices made for the drug-paired substrate (affective bias test) or two pellets-paired substrate (reward learning assay) divided by the total number of trials multiplied by 100 to give a percentage. A value of 50 was then subtracted to give a score where a choice bias towards the drug-paired substrate gave a positive value and a bias towards the control-paired substrate gave a negative value. Choice bias scores, number of omissions and response latency scores during the choice test were analysed using a repeated measures ANOVA with treatment as the within-subject factor and two-tailed paired t-test or S í da k’s multiple comparisons test for post-hoc analysis. Individual positive or negative affective biases were also analysed using a one-sample t-test against a null hypothesis mean of 0% choice bias. For each animal, mean trials to criterion, number of omissions and latency to dig during the affective bias test pairing sessions were analysed using a repeated measures ANOVA with treatment as the factor, and with post-hoc pairwise comparisons made using a two-tailed paired t-test comparison between control (vehicle/low reward:1 pellet) and treatment/manipulation (corticosterone/FG7142/high reward: 2 pellets) for each week (drug-induced negative bias retrieval studies and reward learning assay). Analysis of the choice latency, omissions and trials to criterion was made to determine the presence of any non-specific effects of treatment, such as sedation. Shapiro-Wilk tests were used to determine normal distributions and Mauchly’s sphericity test was used to validate a repeated measures ANOVA. Experimental unit was defined by analysis of the source of greatest variance in the data. For behavioural experiments the experimental unit is animal.

### Electrophysiological data analysis

For acute wash-in experiments, the slice was treated as the experimental unit. For 24-hour post-treatment experiments, a hierarchical approach was used to account for multiple technical replicates per animal. To assess the effect of psilocin on excitatory post-synaptic current (EPSC) amplitude, the mean amplitude during the final 10 minutes of wash-in was calculated as a percentage of the pre-drug baseline. Initial analysis used ordinary least squares (OLS) regression to evaluate the relationship between concentration and response magnitude. Variance heterogeneity across concentrations was assessed using Levene’s test. Gaussian Mixture Modeling (GMM) was then employed to test for the presence of distinct populations. Bayesian Information Criterion (BIC) was used to determine if a single- or two-component Gaussian distribution provided a better fit. Cells were subsequently binned based on GMM clustering. Differences between concentration groups were analyzed using a one-way ANOVA with Tukey’s post-hoc test for pairwise comparisons.

For experiments targeting specific projection populations (CA vs. NT), the change from baseline was determined using paired two-tailed t-tests. Direct comparisons between CA and NT projection groups were performed using Welch’s two-tailed t-test.

24-hour post-acute data were analysed using Linear Mixed-Effects Models (LMM). Treatment and projection (cell-type) were treated as fixed effects, with a Treatment X Projection interaction term included. Animal ID was included as a random effect to account for within-animal correlation. Where a significant interaction or main effect of drug was found, simple effects analysis was performed to compare treatment within each projection level. All simple effects p-values were corrected for multiple comparisons across the three projection types using the Bonferroni method.

## Author contributions

Conceptualisation, E.S.J.R, J.K.H., Z.A.B., and M.D.B.C.; Formal Analysis, J.K.H. and M.D.B.C.; Funding Acquisition, E.S.J.R. and J.R.M.; Investigation, J.B., J.K.H. and M.D.B.C.; Methodology, E.S.J.R., J.K.H and M.D.B.C.; Resources, C.W.T. and G.G.; Supervision, E.S.J.R., C.W.T., G.G., Z.A.B., and J.R.M.; Visualization J.B., J.K.H. and M.D.B.C.; Writing – Original Draft Preparation, J.K.H. and M.D.B.C.; Writing – Review and Editing, J.K.H, M.D.B.C, E.S.J.R, J.R.M., G.G., and Z.A.B.

## Supplementary Materials

**Table S1:** Summary of drug treatments in all animal cohorts.

| Cohort | Treatment | Dose | Route of administration |
| --- | --- | --- | --- |
| 1 | Psilocin | 0, 300 nM, 1 uM | targeted brain infusion |
| 2, 3 | Psilocybin | 0, 0.3, 1.0mg/kg | IP |
| 2, 3 | Volinanserin | 0.1mg/kg | IP |
| 2, 3 | WAY100635 | 0.1mg/kg | IP |
| 1, 2, 3, 4, 5 | FG7142 | 0.0, 3.0mg/kg | SC |
| 4, 5 | DCZ | 0.1mg/kg | PO |

**Table S2A:**
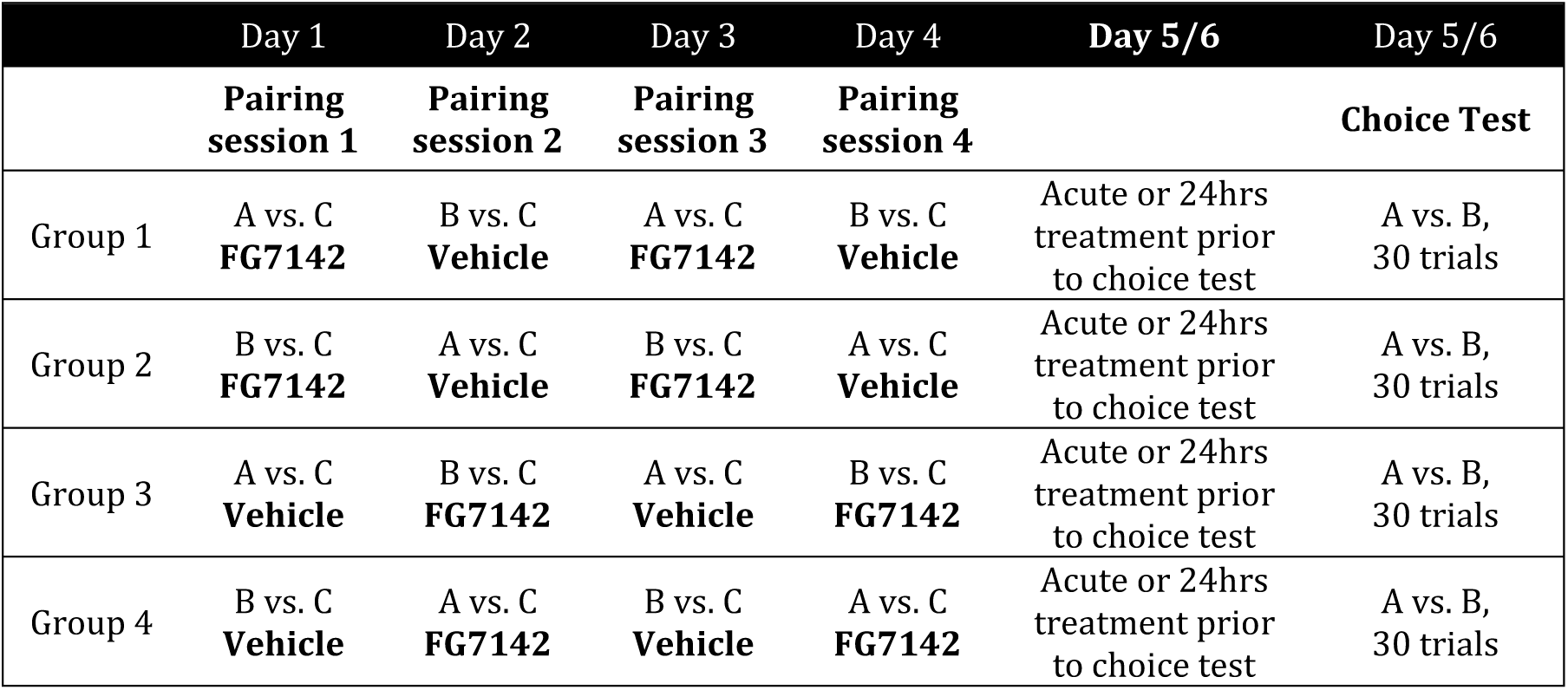
Standard procedure for testing modulation of drug-induced negative affective bias versus vehicle. Each animal receives treatment with FG7142 or vehicle counterbalanced over the four substrate-reward pairing sessions to develop a negative affective state. Substrate (reward-paired substrates - ‘A’ or ‘B’ versus unrewarded substrate - ‘C’) and day are also counter-balanced resulting in four different groups. Prior to choice test each animal receives either vehicle, drug or combination of drugs (vehicle, volinanserin, WAY100635, psilocybin or psilocin).

**Table S2B:** Standard procedure for testing drug-induced affective bias versus vehicle in the new learning studies or reward learning assay. Each animal receives drug treatment/2 pellets or vehicle/1 pellet counterbalanced over the four substrate-reward pairing sessions. Substrate (reward-paired substrates - ‘A’ or ‘B’ versus unrewarded substrate - ‘C’) and day are also counter-balanced resulting in four different groups.

|  | Day 1 | Day 2 | Day 3 | Day 4 | Day 5 |
| --- | --- | --- | --- | --- | --- |
|  | Pairing session 1 | Pairing session 2 | Pairing session 3 | Pairing session 4 | Choice Test |
| Group 1 | A vs. C<br>**Drug/**<br>**2 pellets** | B vs. C<br>**Vehicle/**<br>**1 pellet** | A vs. C<br>**Drug/**<br>**2 pellets** | B vs. C<br>**Vehicle/**<br>**1 pellet** | A vs. B,<br>30 trials |
| Group 2 | B vs. C<br>**Drug/**<br>**2 pellets** | A vs. C<br>**Vehicle/**<br>**1 pellet** | B vs. C<br>**Drug/**<br>**2 pellets** | A vs. C<br>**Vehicle/**<br>**1 pellet** | A vs. B,<br>30 trials |
| Group 3 | A vs. C<br>**Vehicle/**<br>**1 pellet** | B vs. C<br>**Drug/**<br>**2 pellets** | A vs. C<br>**Vehicle/**<br>**1 pellet** | B vs. C<br>**Drug/**<br>**2 pellets** | A vs. B,<br>30 trials |
| Group 4 | B vs. C<br>**Vehicle/**<br>**1 pellet** | A vs. C<br>**Drug/**<br>**2 pellets** | B vs. C<br>**Vehicle/**<br>**1 pellet** | A vs. C<br>**Drug/**<br>**2 pellets** | A vs. B,<br>30 trials |

**Table S3:** Pairing sessions data: number of trials to criterion and latency to dig in the targeted brain infusion studies. Data shown as mean (n=12 animals/group) ± SEM averaged from the two pairing sessions for each substrate-reward association (vehicle or FG7142). There were no significant effects during pairing sessions, either on response latency to dig or number of trials to criterion following treatment with vehicle or FG7142.

| Treatment |  | Response latency (s) |  | Trials to criterion |  |
| --- | --- | --- | --- | --- | --- |
|  |  | Vehicle | FG7142 | Vehicle | FG7142 |
| **Infusion with psilocin 5min. prior to choice test** | Week 1 | 2.0±0.1 | 2.1±0.1 | 6.4±0.1 | 6.4±0.1 |
|  | Week 2 | 1.9±0.1 | 1.9±0.1 | 6.3±0.1 | 6.3±0.1 |
|  | Week 3 | 2.1±0.1 | 1.9±0.1 | 6.2±0.1 | 6.3±0.1 |
| **Infusion with psilocin 24hrs prior to choice test** | Week 1 | 1.7±0.1 | 1.9±0.1 | 6.3±0.1 | 6.3±0.1 |
|  | Week 2 | 1.6±0.1 | 1.5±0.0 | 6.3±0.1 | 6.3±0.1 |

**Table S4:** Choice bias data: response latency to dig in the targeted brain infusion studies. Data shown as mean (n=12 animals/group) ± SEM of an individual latencies during 30 trials of the choice test. No significant difference in latency to make choice was observed in both studies following treatment with vehicle or psilocin.

| Treatment | Dose | Response latency (s) |
| --- | --- | --- |
| Infusion with psilocin 5min.<br>prior to choice test | 0nM | 1.8±0.1 |
|  | 300nM | 1.9±0.1 |
|  | 1000nM | 2.1±0.1 |
| Infusion with psilocin 24hrs<br>prior to choice test | 0uM | 1.5±0.0 |
|  | 300nM | 1.6±0.0 |

**Table S5:** Pairing sessions data: number of trials to criterion and latency to dig. Data shown as mean (n=11-12 animals/group) ± SEM averaged from the two pairing sessions for each substrate-reward association (vehicle or drug/s). There were no significant effects during pairing sessions, either on response latency to dig or number of trials to criterion following treatment with vehicle or any of the drugs. Veh: vehicle, Psi: psilocybin, Vol: volinanserin, Way: WAY100635

| Study |  | Response latency (s) |  | Trials to criterion |  |
| --- | --- | --- | --- | --- | --- |
|  |  | Vehicle | FG7142 | Vehicle | FG7142 |
| Negative bias modulation 24hrs post treatment with vehicle, WAY100635 or volinanserin and vehicle or psilocybin 0.3mg/kg | Week 1 | 2.4±0.1 | 2.6±0.1 | 6.1±0.1 | 6.2±0.1 |
|  | Week 2 | 2.8±0.1 | 2.8±0.1 | 7.0±0.4 | 7.0±0.3 |
|  | Week 3 | 2.7±0.2 | 2.7±0.2 | 7.0±0.2 | 6.6±0.2 |
|  | Week 4 | 3.2±0.3 | 3.2±0.3 | 6.6±0.2 | 6.5±0.2 |
| Negative bias modulation 24hrs post treatment with vehicle or WAY100635 and vehicle or psilocybin 1.0mg/kg | Week 1 | 2.8±0.2 | 2.9±0.2 | 6.6±0.2 | 6.3±0.1 |
|  | Week 2 | 2.6±0.2 | 2.4±0.2 | 6.2±0.1 | 6.3±0.2 |
|  | Week 3 | 2.4±0.2 | 2.5±0.2 | 6.5±0.3 | 6.4±0.2 |
|  | Week 4 | 2.9±0.2 | 3.0±0.2 | 6.3±0.1 | 6.5±0.2 |
| Negative bias modulation 24hrs post treatment with vehicle or volinanserin and vehicle or psilocybin 1.0mg/kg | Week 1 | 2.1±0.1 | 2.1±0.1 | 6.6±0.2 | 6.5±0.2 |
|  | Week 2 | 2.2±0.1 | 1.9±0.1 | 6.9±0.2 | 6.5±0.2 |
|  | Week 3 | 2.3±0.1 | 2.4±0.2 | 6.7±0.2 | 7.1±0.2 |
|  | Week 4 | 2.1±0.1 | 2.1±0.1 | 6.1±0.2 | 6.0±0.2 |
|  |  | Vehicle | Drugs | Vehicle | Drugs |
| New learning with vehicle or WAY100635 and vehicle, psilocybin 0.3mg/kg or psilocybin 1.0mg/kg | Veh/Veh | 2.2±0.1 | 2.4±0.2 | 8.4±0.5 | 7.5±0.4 |
|  | Veh/0.3mg/kg Psi | 2.5±0.3 | 2.4±0.2 | 6.7±0.3 | 7.5±0.5 |
|  | Veh/1mg/kg Psi | 2.6±0.2 | 2.9±0.3 | 7.5±0.5 | 7.0±0.3 |
|  | Way/Veh | 2.5±0.3 | 2.5±0.1 | 7.1±0.4 | 7.6±0.4 |
|  | Way/0.3mg/kg Psi | 2.4±0.2 | 2.6±0.2 | 7.6±0.6 | 7.3±0.5 |
|  | Way/1mg/kg Psi | 2.7±0.2 | 2.9±0.1 | 8.0±0.7 | 7.5±0.6 |
| New learning with vehicle or volinanserin and vehicle, psilocybin 0.3mg/kg or psilocybin 1.0mg/kg | Veh/Veh | 2.3±0.2 | 2.2±0.1 | 7.9±0.5 | 8.0±0.8 |
|  | Veh/0.3mg/kg Psi | 2.6±0.3 | 2.4±0.3 | 6.8±0.3 | 6.9±0.5 |
|  | Veh/1mg/kg Psi | 2.6±0.3 | 2.7±0.4 | 6.8±0.3 | 6.8±0.4 |
|  | Vol/Veh | 2.2±0.1 | 2.6±0.2 | 7.1±0.4 | 7.6±0.5 |
|  | Vol/0.3mg/kg Psi | 2.6±0.2 | 2.3±0.2 | 7.4±0.5 | 7.0±0.2 |
|  | Vol/1mg/kg Psi | 2.4±0.3 | 2.7±0.5 | 7.5±0.5 | 7.5±0.5 |

**Table S6:**
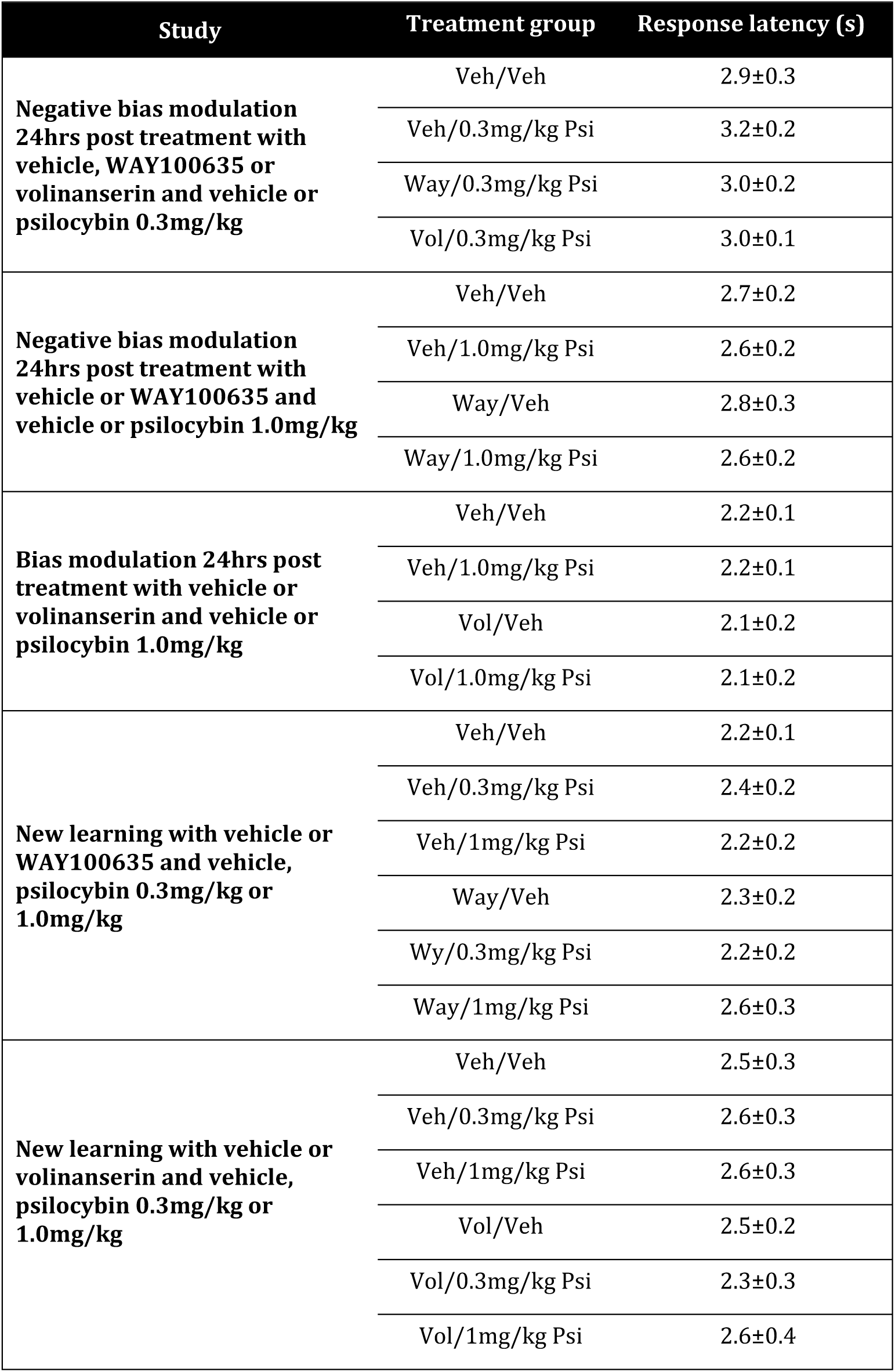
Choice bias data: response latency to dig. Data shown as mean (n=11-12 animals/group) ± SEM of an individual latencies during 30 trials of the choice test. No significant difference in latency to make choice was observed in all studies following treatment with vehicle or any of the drugs. Veh: vehicle, Psi: psilocybin, Vol: volinanserin, Way: WAY100635

| Study | Treatment group | Response latency (s) |
| --- | --- | --- |
| **Negative bias modulation**<br>**24hrs post treatment with**<br>**vehicle, WAY100635 or**<br>**volinanserin and vehicle or**<br>**psilocybin 0.3mg/kg** | Veh/Veh | 2.9±0.3 |
|  | Veh/0.3mg/kg Psi | 3.2±0.2 |
|  | Way/0.3mg/kg Psi | 3.0±0.2 |
|  | Vol/0.3mg/kg Psi | 3.0±0.1 |
| **Negative bias modulation**<br>**24hrs post treatment with**<br>**vehicle or WAY100635 and**<br>**vehicle or psilocybin 1.0mg/kg** | Veh/Veh | 2.7±0.2 |
|  | Veh/1.0mg/kg Psi | 2.6±0.2 |
|  | Way/Veh | 2.8±0.3 |
|  | Way/1.0mg/kg Psi | 2.6±0.2 |
| **Bias modulation 24hrs post**<br>**treatment with vehicle or**<br>**volinanserin and vehicle or**<br>**psilocybin 1.0mg/kg** | Veh/Veh | 2.2±0.1 |
|  | Veh/1.0mg/kg Psi | 2.2±0.1 |
|  | Vol/Veh | 2.1±0.2 |
|  | Vol/1.0mg/kg Psi | 2.1±0.2 |
| **New learning with vehicle or**<br>**WAY100635 and vehicle,**<br>**psilocybin 0.3mg/kg or**<br>**1.0mg/kg** | Veh/Veh | 2.2±0.1 |
|  | Veh/0.3mg/kg Psi | 2.4±0.2 |
|  | Veh/1mg/kg Psi | 2.2±0.2 |
|  | Way/Veh | 2.3±0.2 |
|  | Wy/0.3mg/kg Psi | 2.2±0.2 |
|  | Way/1mg/kg Psi | 2.6±0.3 |
| **New learning with vehicle or**<br>**volinanserin and vehicle,**<br>**psilocybin 0.3mg/kg or**<br>**1.0mg/kg** | Veh/Veh | 2.5±0.3 |
|  | Veh/0.3mg/kg Psi | 2.6±0.3 |
|  | Veh/1mg/kg Psi | 2.6±0.3 |
|  | Vol/Veh | 2.5±0.2 |
|  | Vol/0.3mg/kg Psi | 2.3±0.3 |
|  | Vol/1mg/kg Psi | 2.6±0.4 |

**Table S7:** Pairing sessions data: number of trials to criterion and latency to dig in the chemogenetic manipulation studies. Data shown as mean (n=6^b^-12^a^ animals/group) ± SEM averaged from the two pairing sessions for each substrate-reward association (vehicle or DCZ). There were no significant effects during pairing sessions, either on response latency to dig or number of trials to criterion following these manipulations. DCZ: descoclozapine

| Treatment |  | Response latency (s) |  | Trials to criterion |  |
| --- | --- | --- | --- | --- | --- |
|  |  | Vehicle | FG7142 | Vehicle | FG7142 |
| **DREADDS**<br>**Acute modulation<sup>a</sup>** | Week 1 | 2.1±0.1 | 2.1±0.1 | 6.5±0.2 | 6.3±0.2 |
|  | Week 2 | 1.9±0.1 | 1.8±0.1 | 6.4±0.2 | 6.5±0.2 |
| **DREADDS**<br>**Sustained modulation<sup>a</sup>** | Week 1 | 1.9±0.1 | 1.8±0.1 | 6.3±0.2 | 6.4±0.2 |
|  | Week 2 | 2.0±0.1 | 2.1±0.1 | 6.6±0.3 | 6.2±0.1 |
|  |  | **Vehicle** | **DCZ** | **Vehicle** | **DCZ** |
| **DREADDS**<br>**New learning<sup>b</sup>** | Week 1 | 2.2±0.2 | 2.0±0.1 | 6.2±0.1 | 6.1±0.1 |

**Table S8:** Choice test data: response latency to dig in the chemogenetic manipulation studies. Data shown as mean (n=6^b^-12^a^ animals/group) ± SEM of an individual latencies during 30 trials of the choice test. No significant difference in latency to make choice was observed in all studies following treatment with vehicle or any of the drugs. VEH: vehicle, DCZ: descoclozapine.

| Treatment |  | Response latency (s) |
| --- | --- | --- |
| **DREADDS**<br>**Acute modulation<sup>a</sup>** | VEH | 2.0±0.1 |
|  | DCZ | 1.8±0.1 |
| **DREADDS**<br>**Sustained modulation<sup>a</sup>** | VEH | 1.9±0.1 |
|  | DCZ | 2.0±0.1 |
| **DREADDS**<br>**New learning<sup>b</sup>** | DCZ | 1.8±0.1 |

**Table S9:**
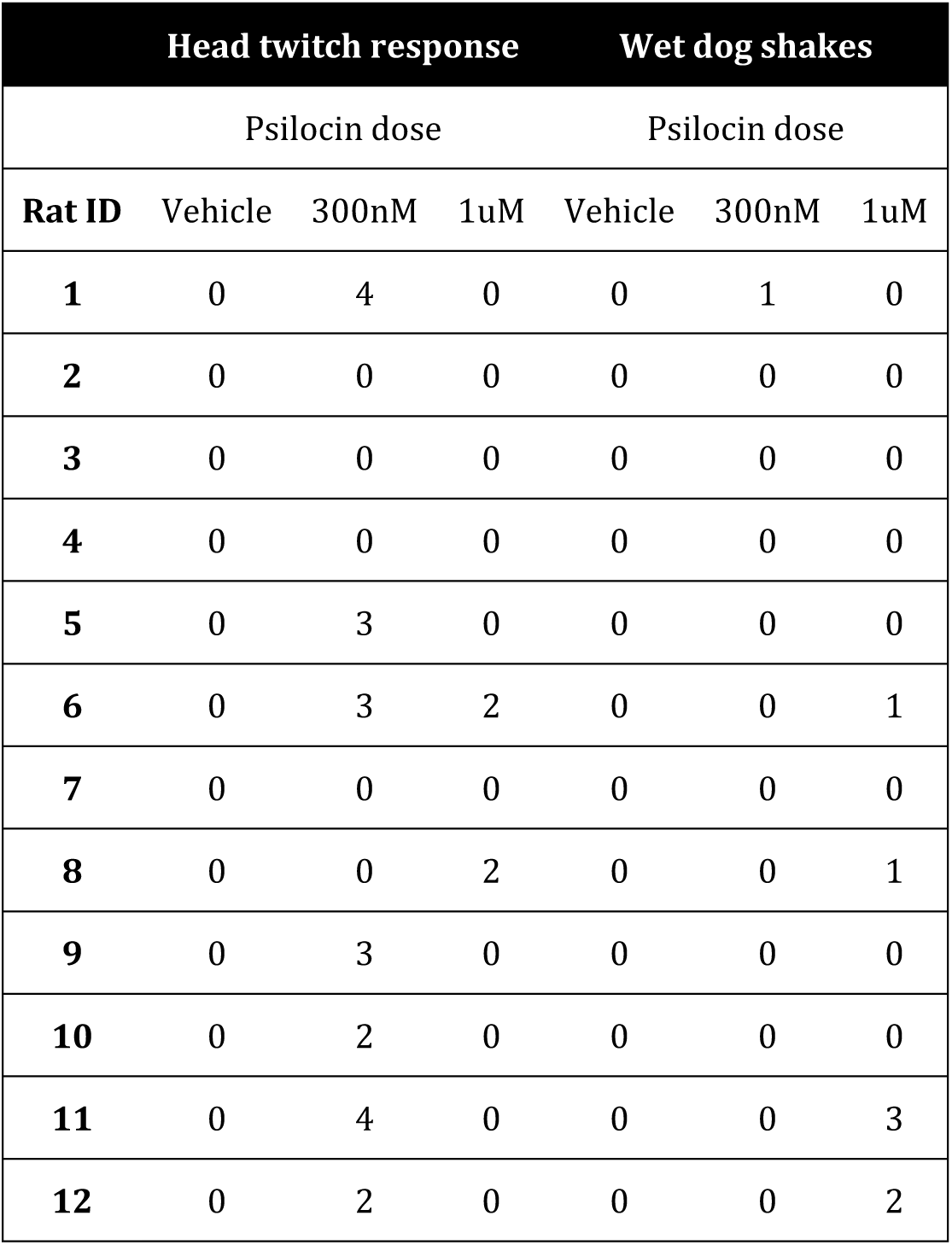
Choice test data: number of head twitches and wet dog shakes following vehicle or psilocin acute infusions. Animals were observed for 15 min. during testing session and the number of head twitches and wet dog shakes were scored.

**Figure S1.**
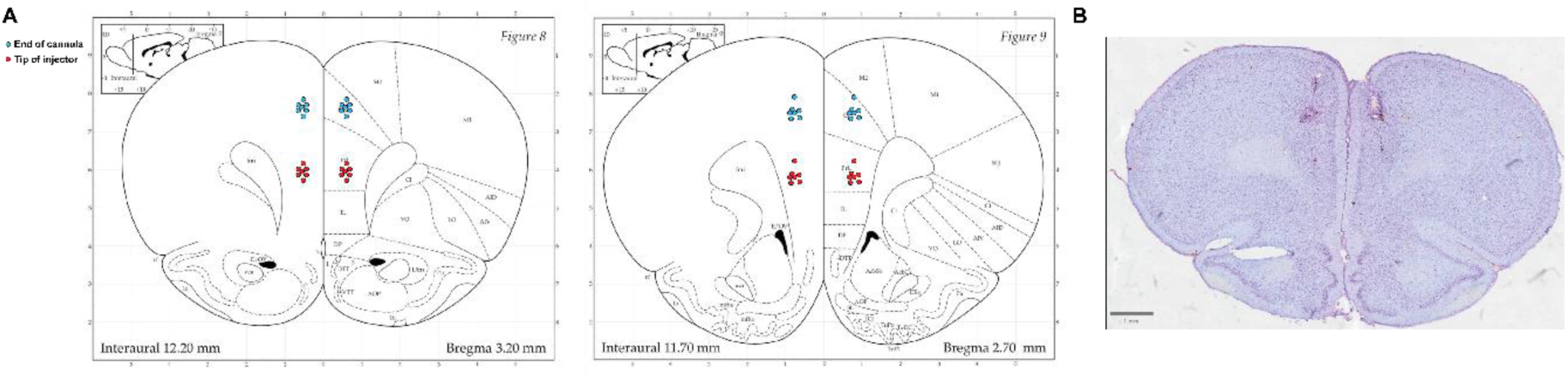
Histological confirmation of prelimbic infusion sites. **A)** Images from rat brain atlas showing histological validation of the end of the cannula and corresponding injector tip sites from the infusion study. **B)** representative image of a cresyl violet stained coronal brain slice from this cohort showing cannulation sites in medial prefrontal cortex.

**Figure S2.**
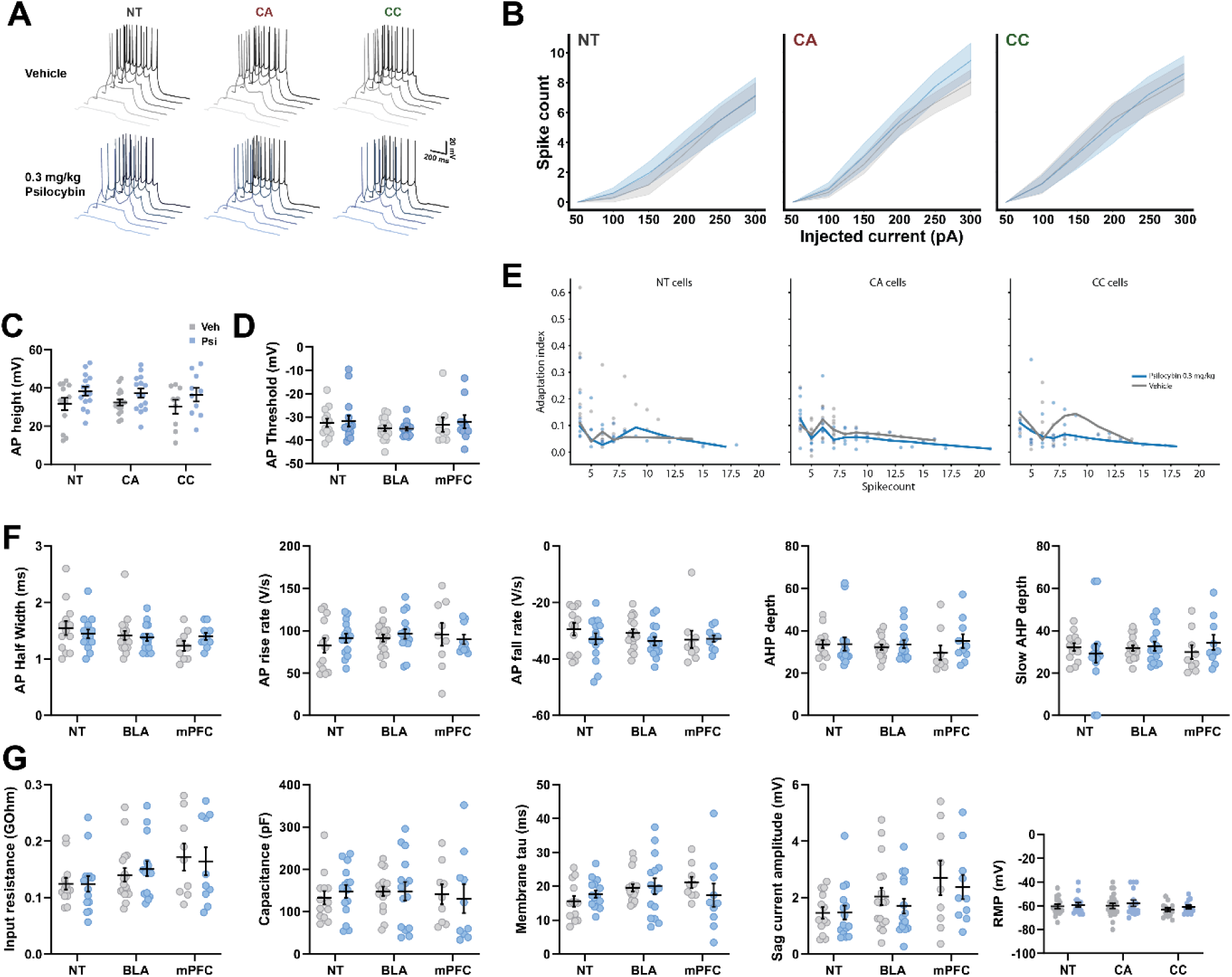
Electrophysiological characterization of prelimbic layer 5 pyramidal cell types following vehicle or 0.3 mg/kg psilocybin. **A)** Representative membrane voltage responses from a series of depolarising current steps (50-300 pA) in non-tagged (NT), cortico-amygdala (CA), and cortico-cortical (CC) cells in slices prepared from Vehicle and Psilocybin (0.3 mg/kg) treated rats. **B)** Spiking vs. injected current relationships in prelimbic cell types averaged across animals. **C-G)** Intrinsic membrane and action potential properties in prelimbic cell-types in vehicle and psilocybin treatment conditions.

## References

1. Goodwin, G.M., et al., Single-dose psilocybin for a treatment-resistant episode of major depression. New England Journal of Medicine, 2022. 387(18): p. 1637–1648.

2. von Rotz, R., et al., Single-dose psilocybin-assisted therapy in major depressive disorder: A placebo-controlled, double-blind, randomised clinical trial. EClinicalMedicine, 2023. 56.

3. Ruhe, H.G., et al., Emotional biases and recurrence in major depressive disorder. Results of 2.5 years follow-up of drug-free cohort vulnerable for recurrence. Frontiers in psychiatry, 2019. 10: p. 145.

4. Hinchcliffe, J.K., et al., Rapid-acting antidepressant drugs modulate affective bias in rats. Science translational medicine, 2024. 16(729): p. eadi2403.

5. Scala, M., et al., Clinical specificity profile for novel rapid acting antidepressant drugs. International Clinical Psychopharmacology, 2023. 38(5): p. 297–328.

6. James, S.L., et al., *Global, regional, and national incidence, prevalence, and years lived with disability for 354 diseases and injuries for 195 countries and territories*, *1990–2017: a systematic analysis for the Global Burden of Disease Study 2017*. The lancet, 2018. 392(10159): p. 1789–1858.

7. Malhi GS, M.J., Depression. Lancet, 2018. 392(10161): p. 24–30.

8. Page, C.E., et al., Beyond the serotonin deficit hypothesis: communicating a neuroplasticity framework of major depressive disorder. Molecular Psychiatry, 2024. 29(12): p. 3802–3813.

9. Cui, L., et al., Major depressive disorder: hypothesis, mechanism, prevention and treatment. Signal transduction and targeted therapy, 2024. 9(1): p. 30.

10. Moncrieff, J., et al., The serotonin theory of depression: a systematic umbrella review of the evidence. Molecular psychiatry, 2023. 28(8): p. 3243–3256.

11. Munkholm, K., A.S. Paludan-Müller, and K. Boesen, Considering the methodological limitations in the evidence base of antidepressants for depression: a reanalysis of a network meta-analysis. BMJ open, 2019. 9(6): p. e024886.

12. Blackburn, T.P., Depressive disorders: Treatment failures and poor prognosis over the last 50 years. Pharmacology research & perspectives, 2019. 7(3): p. e00472.

13. Berman, R.M., et al., Antidepressant effects of ketamine in depressed patients. Biological psychiatry, 2000. 47(4): p. 351–354.

14. Alexander, L., et al., Preclinical models for evaluating psychedelics in the treatment of major depressive disorder. British Journal of Pharmacology, 2024.

15. Robinson, E.S., Delivering a new generation of translational animal models for depression research. Behavioural Pharmacology, 2025. 36(4): p. 175–181.

16. Stuart, S.A., et al., A translational rodent assay of affective biases in depression and antidepressant therapy. Neuropsychopharmacology, 2013. 38(9): p. 1625–1635.

17. Hinchcliffe, J.K., et al., Further validation of the affective bias test for predicting antidepressant and pro-depressant risk: effects of pharmacological and social manipulations in male and female rats. Psychopharmacology, 2017. 234(20): p. 3105–3116.

18. Stuart, S.A., et al., Distinct neuropsychological mechanisms may explain delayed-versus rapid-onset antidepressant efficacy. Neuropsychopharmacology, 2015. 40(9): p. 2165–2174.

19. Shao, L.-X., et al., Psilocybin induces rapid and persistent growth of dendritic spines in frontal cortex in vivo. Neuron, 2021. 109(16): p. 2535–2544.e4.

20. Shao, L.-X., et al., Psilocybin’s lasting action requires pyramidal cell types and 5-HT2A receptors. Nature, 2025: p. 1–10.

21. Vargas, M.V., et al., Psychedelics promote neuroplasticity through the activation of intracellular 5-HT2A receptors. Science, 2023. 379(6633): p. 700–706.

22. Kaufman, J., et al., The 5-HT1A receptor in major depressive disorder. European Neuropsychopharmacology, 2016. 26(3): p. 397–410.

23. Jain, M.K., et al., The polypharmacology of psychedelics reveals multiple targets for potential therapeutics. Neuron, 2025. 113(19): p. 3129–3142.e9.

24. Hesselgrave, N., et al., Harnessing psilocybin: antidepressant-like behavioral and synaptic actions of psilocybin are independent of 5-HT2R activation in mice. Proceedings of the National Academy of Sciences, 2021. 118(17): p. e2022489118.

25. Erkizia-Santamaría, I., et al., *Serotonin 5-HT2A*, *5-HT2c and 5-HT1A receptor involvement in the acute effects of psilocybin in mice. In vitro pharmacological profile and modulation of thermoregulation and head-twich response*. Biomedicine & Pharmacotherapy, 2022. 154: p. 113612.

26. Mengod, G., et al., *Distribution of 5-HT receptors in the central nervous system*, in Handbook of behavioral neuroscience. 2010, Elsevier. p. 123–138.

27. Ly, C., et al., Psychedelics promote structural and functional neural plasticity. Cell reports, 2018. 23(11): p. 3170–3182.

28. Rudebeck, P.H., et al., Prefrontal mechanisms of behavioral flexibility, emotion regulation and value updating. Nature neuroscience, 2013. 16(8): p. 1140–1145.

29. Mertens, L.J., et al., Therapeutic mechanisms of psilocybin: Changes in amygdala and prefrontal functional connectivity during emotional processing after psilocybin for treatment-resistant depression. Journal of Psychopharmacology, 2020. 34(2): p. 167–180.

30. Jiang, Q., et al., Psilocybin triggers an activity-dependent rewiring of large-scale cortical networks. Cell, 2025: p. 2025.08.06.668927.

31. Cameron, L.P., et al., 5-HT2ARs mediate therapeutic behavioral effects of psychedelic tryptamines. ACS chemical neuroscience, 2023. 14(3): p. 351–358.

32. Vollenweider, F.X. and K.H. Preller, Psychedelic drugs: neurobiology and potential for treatment of psychiatric disorders. Nature Reviews Neuroscience, 2020. 21(11): p. 611–624.

33. Savitz, J., I. Lucki, and W.C. Drevets, 5-HT1A receptor function in major depressive disorder. Progress in neurobiology, 2009. 88(1): p. 17–31.

34. Harmer, C.J., R.S. Duman, and P.J. Cowen, How do antidepressants work? New perspectives for refining future treatment approaches. The Lancet Psychiatry, 2017. 4(5): p. 409–418.

35. Likhtik, E., et al., Prefrontal entrainment of amygdala activity signals safety in learned fear and innate anxiety. Nature neuroscience, 2014. 17(1): p. 106–113.

36. Senn, V., et al., Long-range connectivity defines behavioral specificity of amygdala neurons. Neuron, 2014. 81(2): p. 428–437.

37. Adhikari, A., et al., Basomedial amygdala mediates top-down control of anxiety and fear. Nature, 2015. 527(7577): p. 179–185.

38. Poggi, G., F. Klaus, and C.R. Pryce, Pathophysiology in cortico-amygdala circuits and excessive aversion processing: the role of oligodendrocytes and myelination. Brain Communications, 2024. 6(3): p. fcae140.

39. Cai, Y., J. Ge, and Z.Z. Pan, The projection from dorsal medial prefrontal cortex to basolateral amygdala promotes behaviors of negative emotion in rats. Frontiers in Neuroscience, 2024. 18: p. 1331864.

40. Orłowski, P., et al., Naturalistic use of psychedelics is related to emotional reactivity and self-consciousness: The mediating role of ego-dissolution and mystical experiences. Journal of Psychopharmacology, 2022. 36(8): p. 987–1000.

41. Ramos, L. and S.G. Vicente, The effects of psilocybin on cognition and emotional processing in healthy adults and adults with depression: a systematic literature review. Journal of Clinical and Experimental Neuropsychology, 2024. 46(5): p. 393–421.

42. Hamilton, J.P., et al., Depressive rumination, the default-mode network, and the dark matter of clinical neuroscience. Biological psychiatry, 2015. 78(4): p. 224–230.

43. Hamilton, J.P., et al., Default-mode and task-positive network activity in major depressive disorder: implications for adaptive and maladaptive rumination. Biological psychiatry, 2011. 70(4): p. 327–333.

44. Kraehenmann, R., et al., The mixed serotonin receptor agonist psilocybin reduces threat-induced modulation of amygdala connectivity. NeuroImage: Clinical, 2016. 11: p. 53–60.

45. Padawer-Curry, J.A., et al., Psychedelic 5-HT2A receptor agonism alters neurovascular coupling and differentially affects neuronal and hemodynamic measures of brain function. Nature neuroscience, 2025: p. 1–14.

46. Zirkel, R.T., et al., Psilocybin prolongs the neurovascular coupling response in mouse visual cortex. bioRxiv, 2025.

47. Drewko, A.J., R.L. Habets, and T.M. Brunt, Above the threshold, beyond the trip: the role of the 5-HT2A receptor in psychedelic-induced neuroplasticity and antidepressant effects. Molecular Psychiatry, 2025: p. 1–12.

48. Ekins, T.G., et al., Psychedelic neuroplasticity of cortical neurons lacking 5-HT2A receptors. Molecular psychiatry, 2025: p. 1–12.

49. Moliner, R., et al., Psychedelics promote plasticity by directly binding to BDNF receptor TrkB. Nature Neuroscience, 2023. 26(6): p. 1032–1041.

50. Arruda-Carvalho, M. and R.L. Clem, Pathway-selective adjustment of prefrontal-amygdala transmission during fear encoding. Journal of Neuroscience, 2014. 34(47): p. 15601–15609.

51. Saitoh, A., et al., Activation of the prelimbic medial prefrontal cortex induces anxiety-like behaviors via N-methyl-D-aspartate receptor-mediated glutamatergic neurotransmission in mice. Journal of neuroscience research, 2014. 92(8): p. 1044–1053.

52. Suzuki, S., et al., The infralimbic and prelimbic medial prefrontal cortices have differential functions in the expression of anxiety-like behaviors in mice. Behavioural Brain Research, 2016. 304: p. 120–124.

53. Tudi, A., et al., Subregion preference in the long-range connectome of pyramidal neurons in the medial prefrontal cortex. BMC biology, 2024. 22(1): p. 95.

54. Liu, Y., et al., A molecularly defined mPFC-BLA circuit specifically regulates social novelty preference. Science Advances, 2025. 11(17): p. eadt9008.

55. Anastasiades, P.G. and A.G. Carter, Circuit organization of the rodent medial prefrontal cortex. Trends in neurosciences, 2021. 44(7): p. 550–563.

56. Cameron, L.P., et al., A non-hallucinogenic psychedelic analogue with therapeutic potential. Nature, 2021. 589(7842): p. 474–479.

57. Fukumoto, K., et al., Medial PFC AMPA receptor and BDNF signaling are required for the rapid and sustained antidepressant-like effects of 5-HT1A receptor stimulation. Neuropsychopharmacology, 2020. 45(10): p. 1725–1734.

58. Wang, C., et al., Pathway-selective 5-HT1AR agonist as a rapid antidepressant strategy. Cell, 2025.

59. Stuart, S., C.M. Wood, and E. Robinson, Using the affective bias test to predict drug-induced negative affect: implications for drug safety. British journal of pharmacology, 2017. 174(19): p. 3200–3210.

60. Hinchcliffe, J.K. and E.S. Robinson, The affective bias test and reward learning assay: neuropsychological models for depression research and investigating antidepressant treatments in rodents. Current Protocols, 2024. 4(6): p. e1057.

61. Anderson, W.W. and G.L. Collingridge, Capabilities of the WinLTP data acquisition program extending beyond basic LTP experimental functions. Journal of neuroscience methods, 2007. 162(1-2): p. 346–356.

62. Barre, A., et al., Presynaptic serotonin 2A receptors modulate thalamocortical plasticity and associative learning. Proceedings of the National Academy of Sciences, 2016. 113(10): p. E1382–E1391.

63. Berthoux, C., et al., Sustained activation of postsynaptic 5-HT2A receptors gates plasticity at prefrontal cortex synapses. Cerebral Cortex, 2019. 29(4): p. 1659–1669.

